# Genomic Mapping Reveals Cisplatin Disruption of Protein Phosphorylation Signalling Genome-Wide

**DOI:** 10.1101/2024.02.28.582513

**Authors:** Luyu Qi, Qun Luo, Yinzhu Hou, Yan Xu, Wanchen Yu, Xingkai Liu, Bobo Xin, Yaolong Huang, Xiangjun Li, Yanyan Zhang, Shijun Wang, Peter J. Sadler, Yao Zhao, Fuyi Wang

## Abstract

Cisplatin is a DNA-targeting chemotherapeutic. Here we investigate how the cisplatin damaged gene loci are linked to specific protein-driven signalling pathways. Using forward chemical genetics methods, cisplatin damage to specific genes has been mapped in human lung cancer cells, a total of 16216 cisplatin-damaged genes (CDGs) with fold-enrichment > 1.5. Surprisingly, bioinformatics analysis demonstrates that cisplatin targets the majority of human protein kinase and phosphatase genes and involved in 300 core signalling pathways (−log *p* >4). The most associated key signalling pathways are sperm motility and protein kinase A. The highest related disease is cancer, and tissue toxicities related to CDGs are about hepato- and nephro- toxicities. Notably, cisplatin damaged 85% (440) of human protein kinase genes and 81% (110) of human protein phosphatase genes. This suggests that cisplatin acts as a multi-targeting protein-phosphorylation regulator, confirmed by a significant decrease in expression of a series of key protein kinase genes. These results reveal that cisplatin disrupts protein phosphorylation signalling genome-wide.

**Key points summarising the key messages of your article:** A total of 16216 genes damaged by the clinical drug cisplatin in human lung cancer cells were identified using a forward chemical genetics method.

Bioinformatics analysis revealed that cisplatin targets 85% of protein kinase and 81% of phosphatase genes.

Cisplatin acts as a multi-targeting protein-phosphorylation regulator, disrupting protein phosphorylation signalling genome-wide.

The disturbance of protein phosphorylation by cisplatin can be related to anticancer activity and tissue toxicities of the drug.

These findings suggests novel strategies for rational design of next-generation anticancer metallodrugs involving specific targeting of protein phosphorylation.

## Introduction

Cisplatin, *cis*-[PtCl2(NH3)2], is a first-line chemotherapeutic drug, in particular for systemic administration in combination therapy for solid tumours[1,2]. The major target is believed to be nuclear DNA (nDNA)[3,4], although the formation of nDNA adducts is not the sole determinant of cisplatin-induced cytotoxicity[5] and it has been linked to side effects and toxicity[6]. The formation of 1,2-intrastrand -GG-/-AG- (90%) adducts, 1,3-intrastrand -GNG- adducts and interstrand crosslinks unwinds and bends DNA helices, inhibiting DNA replication and transcription, and in turn inducing cell apoptosis and death[3].

The human high mobility group box 1 (HMGB1) protein is well recognized for its specific binding to cisplatin-crosslinked double–stranded DNA [7–10]. HMGB1 functions as a protector of cisplatin-induced DNA damage from nucleotide excision repair[4], and may serve as a negative transcription cofactor to block the binding of transcription factors such as Smad3 to DNA, as revealed by single cell imaging[9]. Shu et al. developed a “cisplatin-seq” method to map cisplatination sites with a single base resolution throughout the whole genome[11]. They revealed that cisplatin was more likely to attack mitochondrial DNA (mDNA) than nDNA, and that the cisplatin damage was enriched within promotors and regions harbouring transcription termination sites[11]. In addition to the cisplatin-seq approach described above, Hu et al. have developed a set of methods, termed “damage-seq” and “eXcision Repair-seq” (XR-seq), to map, at single-nucleotide resolution, the location and repair of cisplatin lesions in genomic DNA adducts[12], respectively. Their mapping data demonstrated that cisplatin damage was uniform and dominated by the underlying sequence, but the repair of cisplatin lesions was heterogeneous throughout the whole genome and affected by multiple factors such as transcription and chromatin states. However, similar to the cisplatin-seq approach, the damage-seq method was not able to assign cisplatin lesions to specific gene loci. To address this issue, the same group integrated their “damage-seq”, “XR-seq” and RNA-seq approaches to study comprehensively the genome-wide profiles of cisplatin lesions in mouse organs[13]. This revealed that the expression of 12 genes was commonly downregulated due to cisplatin damage across mouse kidney, liver, lung, and spleen[13]. However, the combined use of transcriptomics strategies did not produce a full map of cisplatin lesions throughout the whole genome due to the tissue specificity of transcriptomics.

The aim of the research reported here was to map the exact genomic distribution of cisplatin-damaged gene loci by using a forward chemical genetics strategy[14,15]. Here, the HMGB1 box a (HMGB1a)-based affinity microprobes and high throughput sequencing have been combined to screen gene-lesion loci induced by cisplatin on a whole-genome scale (Fig. 1), and to correlate the gene lesions with the action of the drug. Full bioinformatics analysis of cisplatin-damaged genes (CDGs) revealed that cisplatin disrupts the phosphorylation and dephosphorylation of proteins genome-wide. Protein phosphorylation and dephosphorylation are known to be critically important in cancer progression[16].

**Fig. 1.**
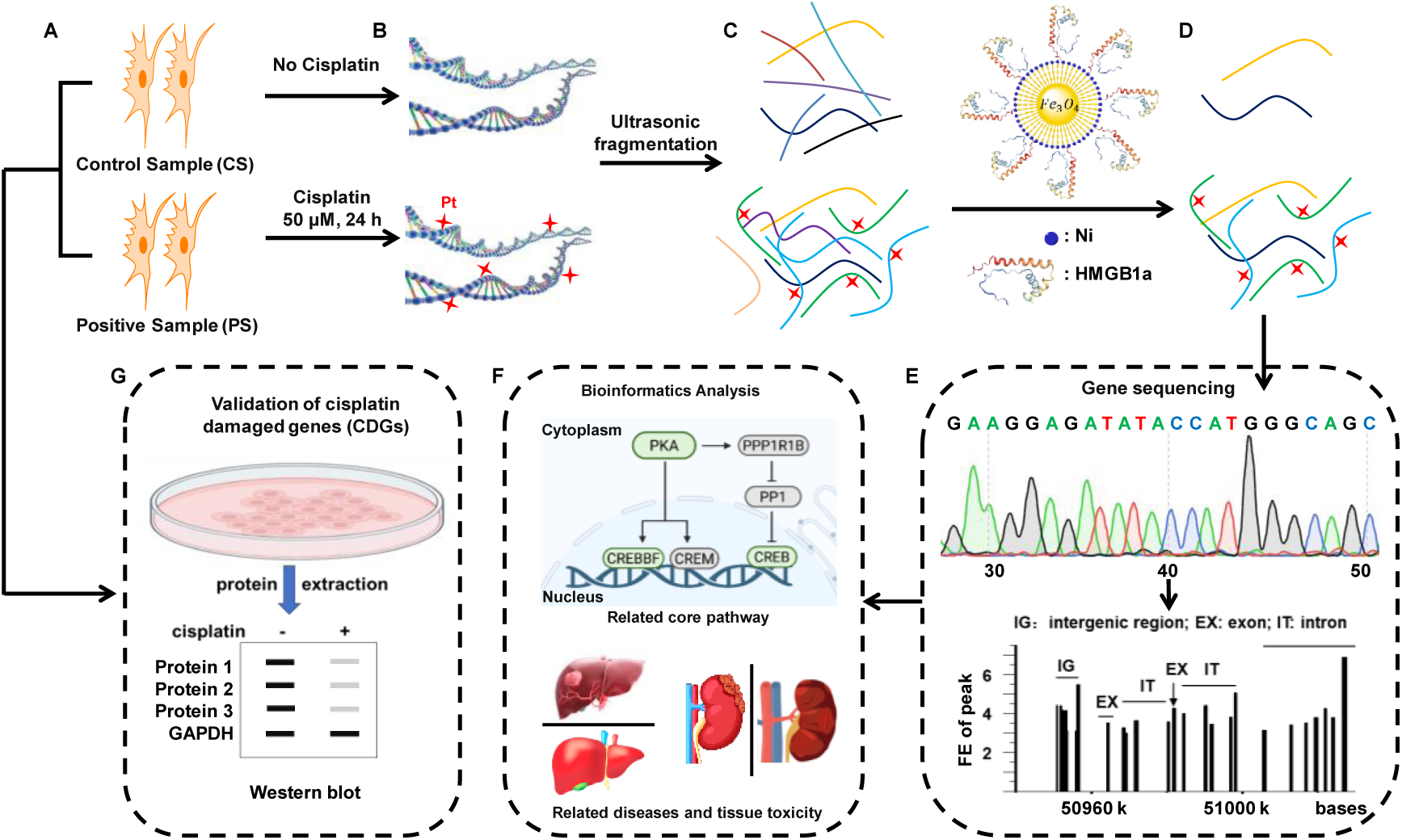
Diagrammatic illustration of the workflow for capture and mapping of gene loci attacked by cisplatin. The procedure includes **(A)** cell culture, **(B)** platination, **(C)** extraction, and fragmentation, **(D)** capture of genomic DNA. **(E)** Sequencing and alignment of cisplatin-damaged DNA fragments. **(F)** Bioinformatics analysis (identification of cisplatin-damaged genes (CDGs)). **(G)** Validation of CDGs by Western Blot assays.

## Results

First an HMGB1a-based affinity probe was constructed to enrich cisplatin-damaged DNA fragments generated by sonication of DNA from human A549 non-small cell lung cancer cells treated with cisplatin. Secondly, cisplatin-damaged DNA fragments were sequenced to map the damaged gene loci. Thirdly, bioinformatics analysis was performed using the Ingenuity Pathway Analysis (IPA) program to correlate cisplatin-damaged genes with core signalling pathways, diseases and toxicity. In addition, the impact of cisplatin damage on the expression of protein-encoded genes was validated using Western Blotting.

### Enrichment and sequencing of cisplatin-damaged DNA

We functionalized Ni-based magnetic microbeads with HMGB1a to assemble affinity microprobes for capturing and isolating 1,2-cisplatin-crosslinked DNA[3,10] from genomic DNA fragments derived from A549 cancer cells treated with 50 μM cisplatin for high throughput Next Generation Sequencing (NGS) (Fig. 1 and Experimental Section). Details of the expression and purification of HMGB1a, construction of affinity microprobes, cell harvesting, cisplatin-damaged DNA extraction, fragmentation and sequencing are described in the Experimental Section in the Supplemental Materials and summarized in Fig. 1.

Since HMGB1 binds to cisplatin-crosslinked DNA with high affinity, but can also bind to bent native DNA[3], we extracted and fragmented genomic DNA from A549 cells with (positive samples, PS) and without (control samples, CS) exposure to cisplatin for DNA sequencing. These DNA fragments were then subjected to the high-throughput sequencing after the determination of the platination level (Fig. 2A) and the quality check (Fig. 2B and 2C).

**Fig. 2.**
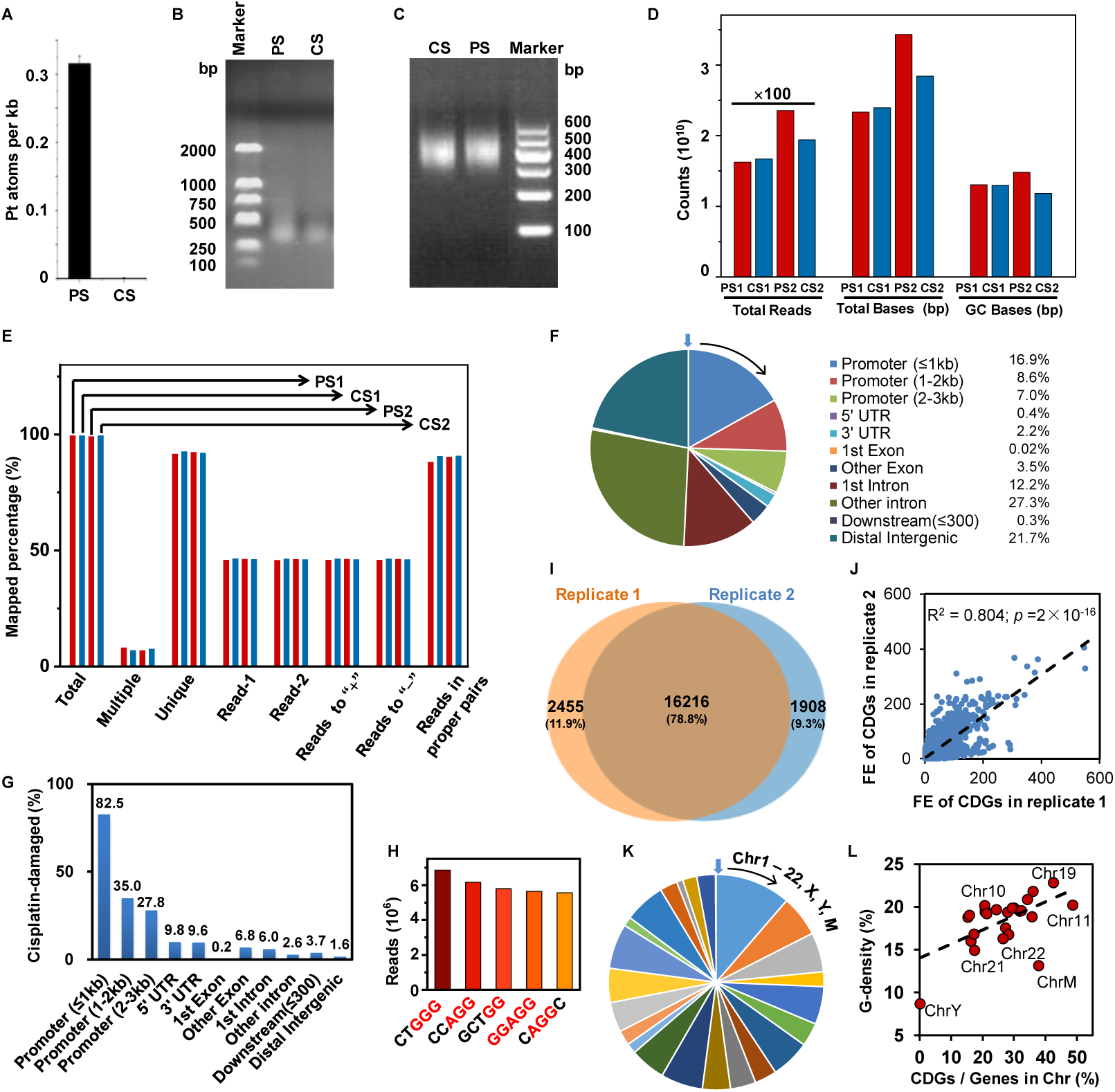
Mapping of cisplatin-damaged genes. **(A)** The platination level of DNA fragments extracted from positive sample (PS) and control sample (CS), determined by ICP-MS. **(B)** The size distribution of DNA fragments from PS and CS groups. **(C)** The size distribution of sequences in DNA libraries constructed from captured DNA fragments. **(D)** Counts of total read, total bases and GC bases mapped to PS and CS groups. **(E)** Percentage of mapped genes for PS and CS groups. Total: total mapped genes; multiple: multiple mapped genes; unique: unique mapped genes; read-1 & 2: first and second read; reads to “+” & “−”: read from 5′- to 3′-end (+) and 3′- to 5′-end (−); reads in proper pairs: reads with proper base-pair matching. **(F)** Distribution of cisplatin-damaged DNA fragments in genomic regions. **(G)** Percentage of the lengths of the cisplatin-damaged DNA fragments in specific genomic regions. **(H)** The five pentanucleotide motifs that were the most frequently detected in CDGs. **(I)** Venn Diagram of total CDGs with FEG >1.5 in two independent biological replicates. **(J)** Correlation analysis of the normalized FEG of CDGs in two replicates. **(K)** Distribution of CDGs over 23 chromosomes and mitochondria (Extended Table 3). **(L)** Correlation of the relative G-density in the chromosome (y-axis) with the proportion of CDGs detected in a chromosome to total number of genes in the human genome (x-axis).

We minimize the risk of false negative results from gene-mapping, as described in the Experimental Section in the Supplemental Materials (Sequencing and gene-mapping). The total read counts and total base counts of the positive samples (PS) did not show a pronounced difference from those of the control samples (CS) (Fig. 2D). However, because cisplatin prefers to crosslink -GG-sites on DNA, the GC base-pair counts of the PS accounted for a slightly higher ratio to the total base counts than those of the CS (Fig. 2D; Supplementary Table S1). Consequently, the mapped percentage of total reads in properly-matched base pairs of positive samples were slightly lower than those of control samples (Fig. 2E; Supplementary Table S1). Nevertheless, there are no significant differences in the total mapping percentage (≈ 99%) and unique mapping percentage (> 91%) of genomic DNA fragments from both PS and CS. These data show that the quality of the sequencing is reliable to characterize cisplatin damaged genes (CDGs) throughout the whole genome.

### Mapping of cisplatin-damaged genes

To identify cisplatin-damaged genes (CDGs), we compared the total reads of all peaks mapped to a gene in the positive sample (PS) with those of all peaks mapped to the same gene in the corresponding control sample (CS). If the ratio [number of peaks]PS/[number of peaks]CS, the fold-enrichment of a peak (FEP) mapped to a gene, was ≥ 1.5, the peak was counted to the gene[17]. The fold-enrichment of a gene (FEG) is the sum of the FEP of all peaks mapped to the gene (details in Materials and Methods in the Supplementary Materials). Notably, to eliminate the bias arising from the different total reads of all peaks mapped in all samples, the normalized value of the read of a peak to the total read of all peaks for the sample were used (Supplementary Table S1). Following this method, we mapped 246,424 and 362,594 peaks with FEP ≥ 1.5 from replicates 1 and 2, respectively. As shown in Fig. 2F, the peaks mapped to promoter regions, located 3k base pairs upstream of a gene, account for the highest percentage (32.48%) of all peaks mapped genome-wide. Considering the different lengths of each region in the genome, the coverage of peaks mapped to the promotor regions of all CDGs account for 82.5% of the total length of promotor region 1 (≤ 1 kb), 35.0% of promotor region 2 (1 – kb), 27.8% of promotor region 3 (2 – 3 kb) (Fig. 2G). These coverages are much higher than for peaks mapped to other regions, e.g. exon and intron regions. These results show that cisplatin damage is enriched in the promotor regions, in particular, in regions close to encoding regions, in line with previous reports[11]. Since HMGB1a binds to double-stranded DNA over 5 base-pairs[3], the occurrence frequency of pentanucleotide motifs for all peaks containing platination sites was calculated, showing that the top 5 most-frequently platinated pentanucleotide motifs are CTGGG, CCAGG, GCTGG, GGAGG and CAGGC (Fig. 2H). This is consistent with the well-established finding that cisplatin prefers to bind to GG sites on DNA[3,4].

When the FE of a CDG was calculated, the peaks mapped to the distal intergenic region of a gene were discarded. Following this procedure, a total of 16216 CDGs with FEG ≥1.5 were mapped which were common in the two replicates (Fig. 2I; Supplementary Table S2). Moreover, the FEG of CDGs mapped in the two replicates were linearly correlated, with R = 0.804, as shown in Fig. 2J, evidence of good reproducibility of the sequencing. The four CDGs with the highest FEG mapped in this way are shown in Supplementary Fig. S1.

The 16216 CDGs are unevenly distributed over all 23 pairs of chromosomes (Fig. 2K and Supplementary Table S3), showing a moderate dependence on the average G-density of the chromosomes, i.e., the higher the average G-density of a chromosome, the more the genes are attacked by cisplatin (Fig. 2L and Supplementary Table S3), again consistent with the known preference of cisplatin for crosslinking -GG-sites.

### Bioinformatics Analysis

The Ingenuity Pathway Analysis (IPA) program was employed to perform genome-wide annotation on the mapped CDGs. The 16216 CDGs were input into the IPA data pool and matched to 14971 protein-encoding genes in the human genome (hg19), including 2615 enzymes, 1488 transcription regulators, 810 transporters, 634 kinases and 205 phosphatases (Fig. 3A, Supplementary Table S4). Eleven microRNA- (miRNA) encoding genes were also identified, through these genes have a relatively low FEG as their sequences are much shorter than genomic DNA.

**Fig. 3.**
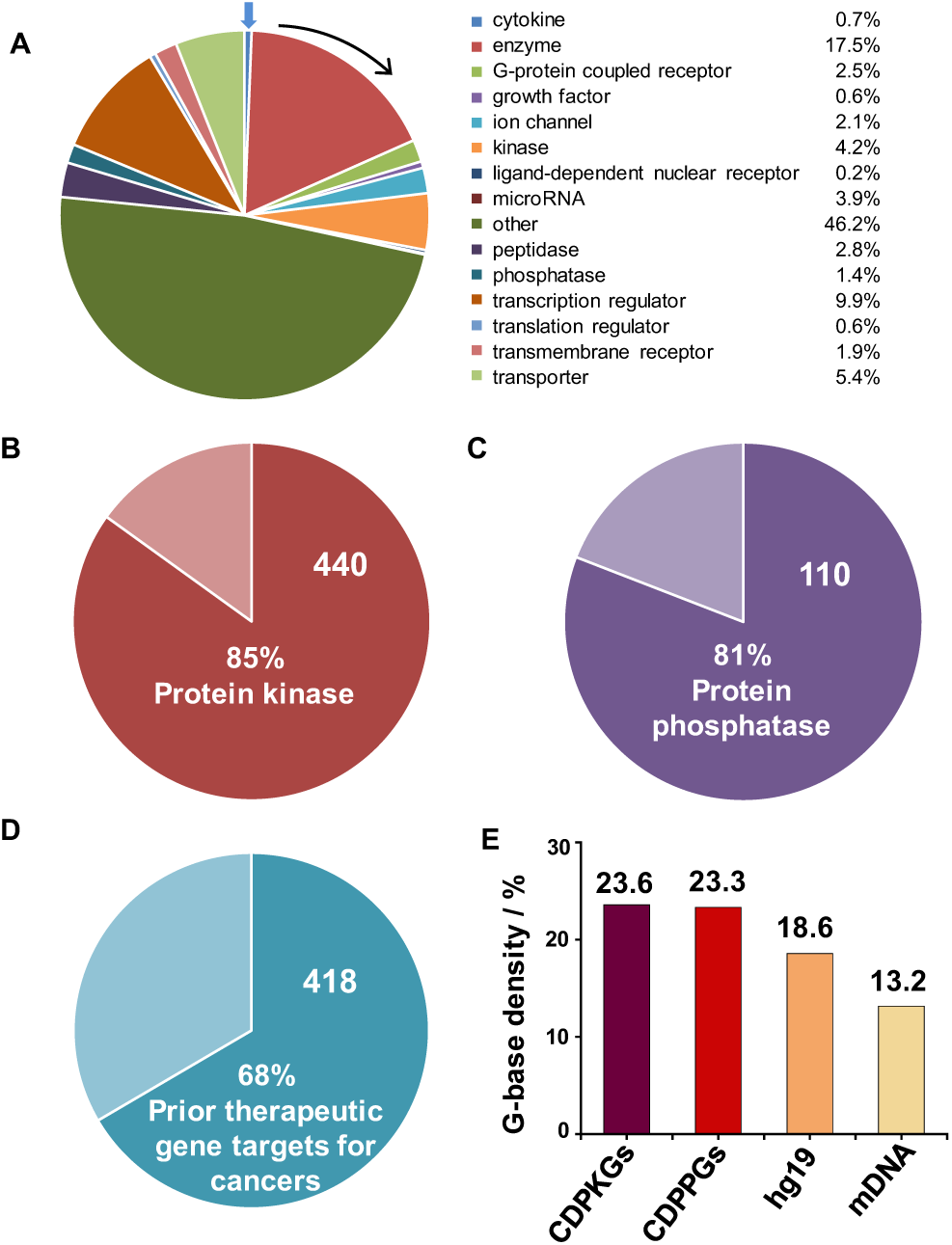
**(A)** Biological function classification of 14971 protein-encoding cisplatin-damaged genes (CDGs) with FEG >1.5. **(B)** Ratio of cisplatin-damaged protein kinase genes (CDPKGs) in all the 518 reported putative protein kinase genes in the human genome[18]. **(C)** Ratio of cisplatin-damaged protein phosphatase genes (CDPPGs) in all the 136 reported protein phosphatases in the human genome[19]. **(D)** Ratio of CDGs in the 628 therapeutic gene targets for cancers identified by the CRISPR/Cas9-based screening[20]. **(E)** Average percentage of G bases in CDPKGs and CDPPGs, hg19 and mDNA.

Notably, among the 634 cisplatin-damaged kinase genes, 440 are protein kinase genes (Supplementary Table S5), accounting for 85% of 518 reported putative protein kinase genes in the human genome (Fig. 3B)[18]. There are also 110 protein phosphatase genes (CDPPGs) among the 205 cisplatin-damaged phosphatase genes (Supplementary Table S6), accounting for 81% of the 136 reported protein phosphatases in the human genome (Fig. 3C)[19]. Significantly, among the CDGs, 418 genes belong to the 628 prior therapeutic gene targets for cancers identified previously by the CRISPR/Cas9-based screening (Fig. 3D), 54 (13%) of which are targets of clinically-used drugs or candidates in preclinical development[20] (Supplementary Table S7). Also, the average percentages of G-bases in CDPKGs and CDPPGs are significantly higher than that of all the genes in the human genome (hg19, Fig. 3E). Therefore, platination on the whole genome is preferential for genes with high G-density, consistent with previous reports[3,4,11]. For lung cancer cells, only 14 cisplatin-damaged mitochondrial genes were identified with low FEG ranging from 1.5 to 2.4 (Supplementary Table S8). Considering the much lower G-density in mDNA (Fig. 3E), it is reasonable that mitochondrial DNA has a low platination level.

It is notable that among the 16216 CDGs with FEG >1.5, there are 36 pseudogenes (Supplementary Table S2). In general, pseudogenes are thought to be non-functional “junk genes”. However, increasing evidence argues that pseudogenes can act as key regulators at DNA, RNA or protein levels in various human diseases such as cancer[21,22].

### Highly associated core signalling pathways of cisplatin-damaged genes

Cisplatin-damaged genes with FEG > 20 (4774 genes) were selected and processed by IPA. They are associated with 300 core signalling pathways (CSPs) with –log(*p*) > 4, i.e. *p*<0.0001 (Supplementary Table S9). The top most associated pathways of these CDGs includes RHO GTPase cycle, netrin, synaptogenesis, myelination, glutaminergic receptor (enhanced), sperm motility, actin cytoskeleton, integrin, Rho family GTPases, and protein kinase A signalling pathways (Fig. 4A). The detailed information of these signalling pathways is in the Supplementary Figs. S2 – S18.

**Fig. 4.**
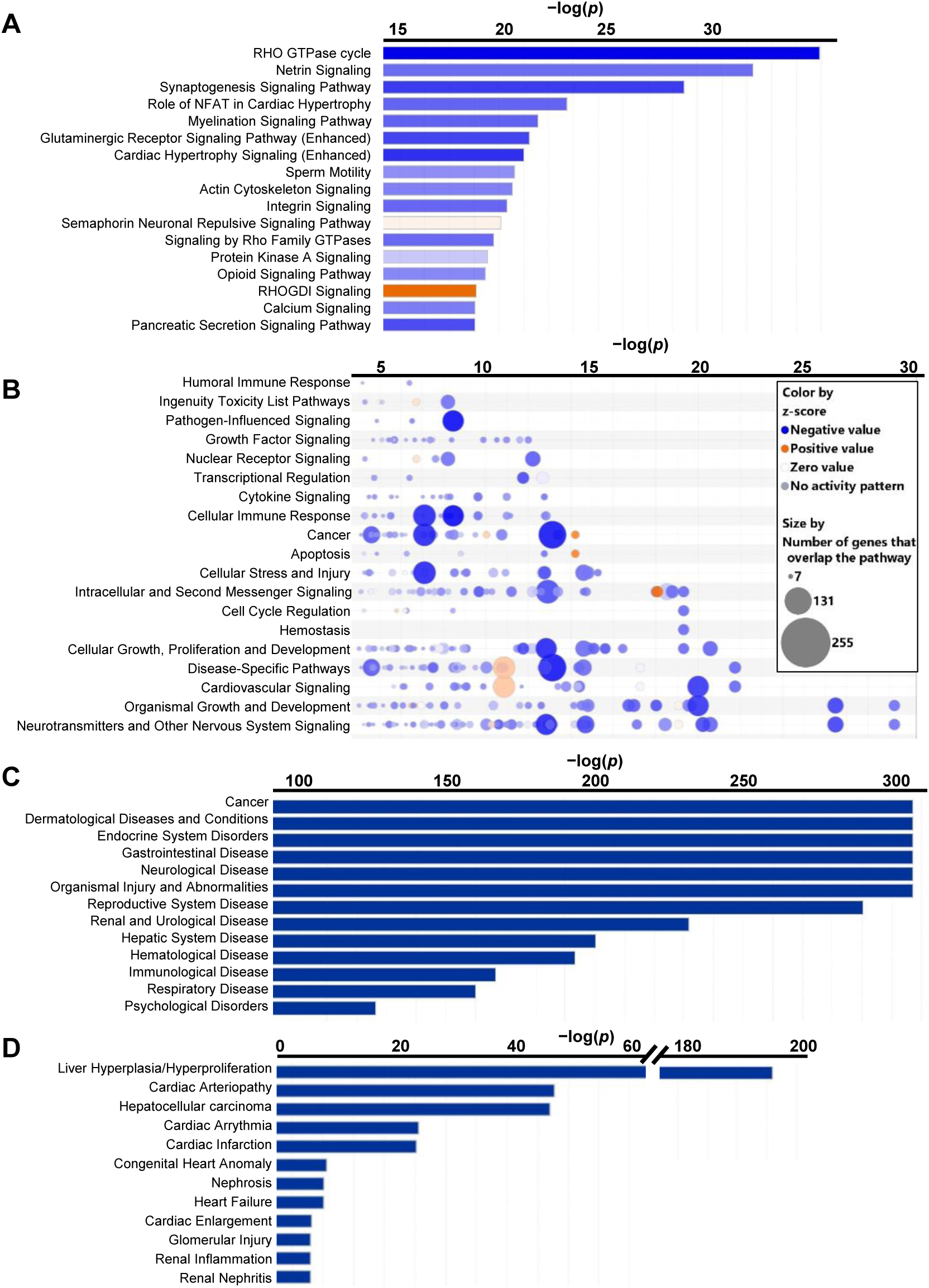
Bioinformatics analysis of cisplatin-damaged genes (CDGs). **(A)** The core signalling pathways (CSPs) which cisplatin-damaged genes (CDGs) are associated with. **(B)** The CDGs-associated CSPs can be further classified into 19 categories. **(C)** Diseases and functions which CDGs are associated with. **(D)** Toxicity which CDGs are associated with.

All signalling pathways associated with the CDGs can be classified into 19 categories (Fig. 4B). The highest associated categories (with negative z-scores) such as organismal growth and development, cellular growth, proliferation and development, cell cycle regulation, intracellular and second messenger signalling; cellular stress and injury, apoptosis, cancer, and cellular immune response, are closely related to the inhibition of cancer cell growth. Also, those categories such as neurotransmitters and other nervous system signalling pathways, and cardiovascular signalling pathways, may be closely related to the known side effects of cisplatin, such as neurotoxicity and cardiotoxicity [6,23].

Furthermore, IPA revealed that among the top 300 CDGs associated CSPs (−log *p* <4), cisplatin-damaged protein kinase genes (CDPKGs) on average account for 24 % of the genes in each CSP (Supplementary Table S9). This proportion is much higher than that (4.8%) of CDPKGs to the 4774 CDGs with FEG >20 in the whole genome (Supplementary Table S5). This suggests that the damage on protein kinase genes plays a significant role in the mechanism of action of cisplatin.

It is notable that CDGs are highly associated with protein kinase A (PKA) signalling pathway with −log *p* = 18.5 (Fig. 4A, Supplementary Fig. S12), and 152 CDGs including 28 (18.4%) CDPKGs, are involved in this signalling pathway (Supplementary Table S9 and S11). Among the CDPKGs involved in PKA signalling pathway, a series of protein kinases showed high fold-enrichment (FEG>100), e.g. *PRKAG2*: 251, *PRKCE*: 161, *PRKCB*: 133, *PRKCZ*: 115, and *PRKAR1B*: 112, which indicates that protein kinases play crucial roles in the activity of cisplatin, meriting further investigation as a potential targets for cancer treatment.

Another core signalling pathway with which the CDGs are highly associated is the molecular mechanisms of cancer signalling pathway (−log *p* = 13.1) (Supplementary Table S9, Supplementary Fig. S16). 255 CDGs are involved in this pathway, and cover 66% of all genes in it. The very negative *z* score (−13.2) predicted by IPA demonstrated that these gene lesions induced by cisplatin downregulate the development of cancers. This is consistent with the clinical therapeutic effect of cisplatin.

We found that CDGs are also highly associated with the non-small cell lung cancer (NSCLC) signalling pathway (−log *p* = 9.5) (Supplementary Table S9, Supplementary Fig. S18). Forty-seven genes, including 12 PKGs, of this core signalling pathway are damaged by cisplatin, covering 63% of all the genes in it. The *z* score of this pathway is −4.4, which therefore predicts to be significantly inhibited by cisplatin. Many patients with non-small-cell lung cancer have no response to the tyrosine kinase inhibitor gefitinib, which targets the epidermal growth factor receptor (EGFR)[24]. However, cisplatin can damage *EGFR* gene in the NSCLC signalling pathway (Supplementary Table S2 and protein expression data shown in Fig. 5), which may play a role in the anticancer activity of cisplatin towards NSCLC. Given that AKT/PKB can promote survival of NSCLC[25], damage to the *AKT* gene by cisplatin may contribute to the killing of the NSCLC cancer cells.

**Fig. 5.**
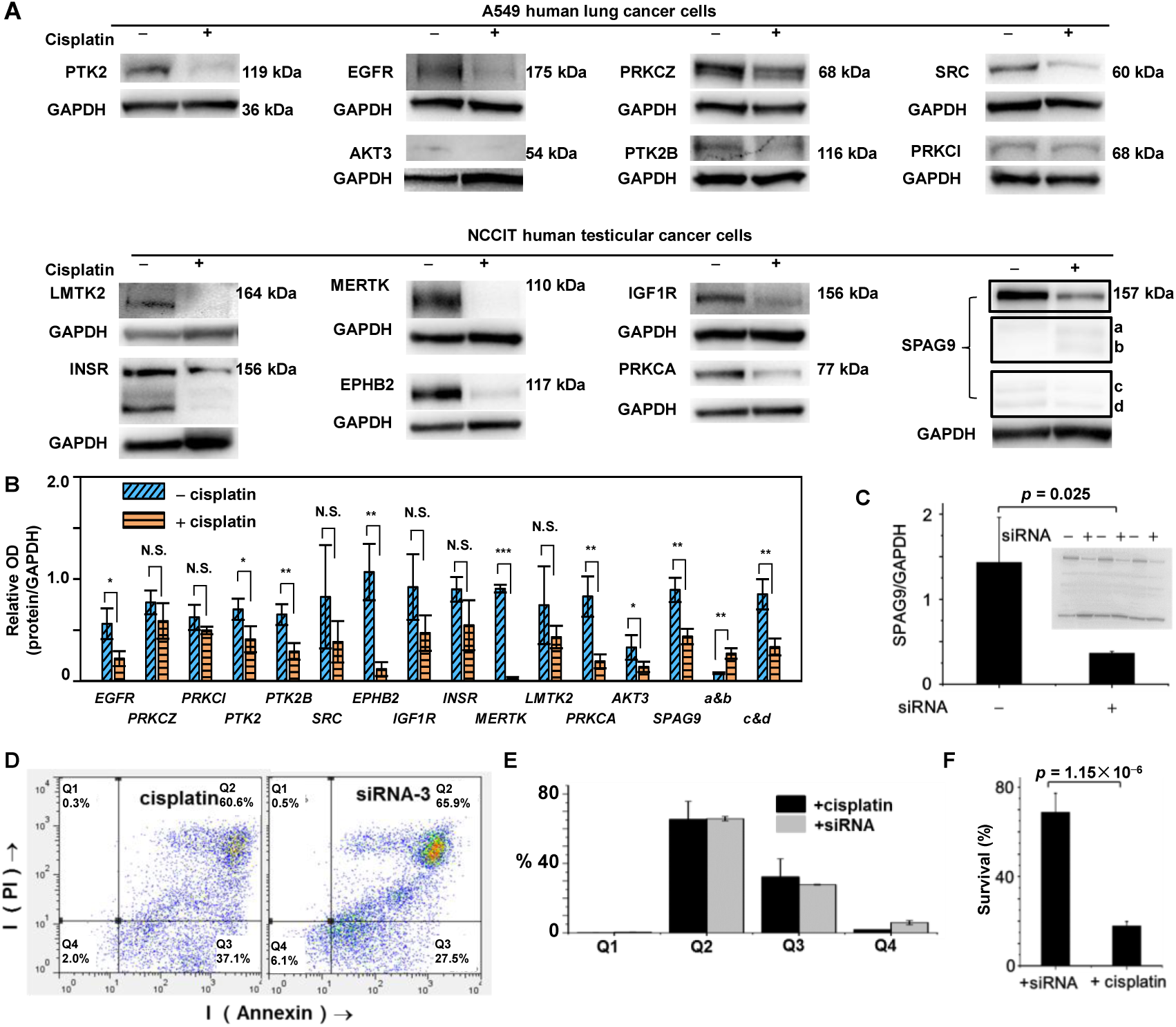
Western Blot assays and siRNA silencing. **(A)** Western Blot images of selected gene-encoding proteins expressed in A549 lung and NCCIT testicular cancer cells with (+) and without (−) cisplatin (12 μM) treatment. GAPDH (Glyceraldehyde-3-phosphate dehydrogenase) was the internal reference. The a – d subunits of JIP-4 (expressed by *SPAG9*) are shown in Supplementary Fig. S20A. **(B)** Optical density (OD) ratio of a gene-encoded protein to GAPDH. **(C)** Western Blot data for siRNA silencing of *SPAG9*. **(D)** Flow cytometric analysis of viability (Q1), early-stage apoptosis (Q2), late-stage apoptosis (Q3) and necrosis (Q4) for NCCIT cells subjected to cisplatin treatment in comparison with siRNA silencing of *SPAG9*. **(E)** The quantitative analysis of the flow cytometric analysis shown in D. **(F)** Survival rate (%) of NCCIT cells subjected to cisplatin treatment or siRNA silencing of *SPAG9*. In B, two-tailed unpaired Student′s t-test was used to statistics process of the Western Blot assay (n = 3). N.S.: no significant; * indicates *p* < 0.05, ** *p* < 0.01, *** *p* < 0.001. *pEGFR*=0.0244, *pPRKCZ*=0.2057, *pPRKCI*=0.1615, *pPTK2*=0.0366, *pPTK2B*=0.0088, *pSRC*=0.2343, *pEPHB2*=0.0044, *pIGF1R*=0.1000, *pINSR*=0.0893, *pMERTK*=2.71×10^-6^, *pLMTK2*=0.2453, *pPRKCA*=0.0062, *pAKT3*=0.0126, *pSPAG9*-full=0.0049, *pSPAG9*-a&b=0.0030, *pSPAG9*-c&d=0.0061.

We also found IL-15 production and sperm motility (SP) signalling pathways, and a series of neural system and cardiac related signalling pathways. The details are shown in Supplementary Tables S9 and S10 and Supplementary Figs. S2 – S11, S13 – 15, S17. The disruption to these signalling pathways by cisplatin may be related to the side effects of this drug.

### Association of cisplatin-damaged genes with diseases and tissue toxicities

The association of CDGs with diseases and functions were analysed by IPA and the most associated disease of CDGs is cancer (−log *p* > 300) (Fig. 4C). The top five types of cancers with negative z-scores (predicting inhibition) to which CDGs are highly related are development of carcinoma (*z*-score −2.3), nonhematologic malignant neoplasm (*z*-score −3.8), abdominal carcinoma (*z*-score −3.6), abdominal adenocarcinoma (*z*-score −2.3), digestive system cancer (*z*-score −3.9), (Supplementary Table S12, Supplementary Fig. S19). The negative z-scores for these cancers predict that they may be associated with cisplatin-damaged genes.

It is notable that the sperm-associated antigen 9 (*SPAG9*) gene is involved in 147 of 361 cancers or functions with which the CDGs are highly associated (Supplementary Table 13). *SPAG9* abundantly expresses C-jun-amino-terminal kinase-interacting protein 4 (JIP-4) which consists of four subunits (Supplementary Fig. S20A) in testicular haploid germ cells and is essential for the development, migration and invasion of cancer[26]. JIP-4 as a scaffolding protein knits mitogen-activated protein kinases (MAPKs) and their transcription factors to activate specific phosphorylation signalling pathways, such as MAPK and PI3K (phosphatidylinositol 3’-kinase) pathways.[27] These imply that cisplatin damage on *SPAG9* (FEG = 62.0) may play a crucial role in the anticancer activity of cisplatin (confirmed by gene silencing and cell apoptosis experiments, Fig. 5C – F).

Tissue toxicity is a major factor which limits the clinical application of cisplatin. However, the molecular basis for its toxicity remains unclear. IPA annotation of the associated toxicity of cisplatin damage to genomic DNA suggests that the CDGs are closely associated with liver, heart, and kidney (Fig. 4D). The most highly related tissue toxicities are liver hyperplasia or hyperproliferation (−log *p* 195.9), cardiac arteriopathy(−log *p* 47.7), hepatocellular carcinoma (−log *p* 47.0), cardiac arrhythmia (−log *p* 25.6), cardiac Infarction (−log *p* 25.2), congenital heart anomaly (−log *p* = 10.6), nephrosis (−log *p* 10.1), heart failure (−log *p* 10.1), cardiac enlargement (−log *p* 8.1), glomerular injury (−log *p* 7.9), renal inflammation (−log *p* 7.9), and renal nephritis (−log *p* 7.9) (Fig. 4D and Supplementary Table S14). Since liver hyperplasia or hyperproliferation are usually related to liver tumours, liver carcinoma and liver cancer, and the z-scores are negative (−2.5, −2.9, and −3.4 respectively), this result suggest that cisplatin-damaged genes may be related to therapeutic effects. Also, as reported, cisplatin accumulate mostly in liver and kidney rather than in heart, the most relevant toxicity to all CDGs is likely nephrotoxicity [6,23,28]. However, further investigation is needed to correlate the gene lesions induced by cisplatin with the phenotypes of clinical nephrotoxicity.

### Protein expression in cisplatin-treated cancer cells

To investigate the relationship between the CDGs and protein expression, classical Western Blot assays were performed. The expression level of 15 protein kinase genes involved in sperm mobility signalling pathway, with which the CDGs are highly associated (Fig. 4A), was studied with or without cisplatin exposure (details in Supplementary Table S15). *SPAG9* (sperm associated antigen 9) was also chosen because it expresses JIP-4 protein which activate a series of phosphorylation signalling pathways[27]. Since these chosen genes are expressed at different levels in different types of cells, we selected two human cancer cell lines, A549 (non-small cell lung cancer cell line) and NCCIT (testicular cancer cell line) to perform Western Blot assays. Testicular cancer is curable by cisplatin.

As shown in Fig. 5A, the expression levels of 14 selected genes decreased after cisplatin treatment. The changes in expression levels of 8 genes, including 7 protein kinase genes and *SPAG9*, were significant (*p* < 0.05) or very significant (*p* < 0.01) (Fig. 5B). The data indicate that these genes were indeed damaged by cisplatin, substantiating the gene-mapping results described above. However, there is no direct correlation between the FEG of a specific CDG, or the base-coverage of cisplatin-damaged peaks over a damaged gene and the fold-change in expression of the gene (Supplementary Fig. S21). This is probably attributable to the complexity of the gene expression machinery (Supplementary Fig. S20B).

Given that *SPAG9* specifically expresses JIP-4 in high abundance in testicular haploid germ cells, its cisplatin-damage may account for the high cure rate of cisplatin for testicular cancer. Indeed, we found that siRNA silencing of the *SPAG9* gene (Fig. 5C) induced a similar level of early- and late-stage apoptosis in NCCIT cells subjected to cisplatin treatment (Fig. 5D – F). This demonstrated that *SPAG9* plays an important role in the action of cisplatin as the upstream mediator of phosphorylation pathways (Supplementary Fig. S22A and B).

The expression of the receptor tyrosine kinase *MERTK* (c-mer proto-oncogene tyrosine kinase) was the most strongly inhibited (fold change = 32) due to cisplatin damage (Fig. 5A, B). This receptor phosphorylates Akt1-Y26, mediating Akt activation and survival signalling, which in turn drives oncogenesis and therapeutic resistance [29]. Thus, cisplatin damage to *MERTK* may be implicated in the mechanism of action of cisplatin. Hence, *MERTK* is also a potential therapeutic target [29,30].

The expression of *MERTK*-related gene *AKT3* (RAC-gamma serine/threonine-protein kinase) was inhibited 3.5-fold by cisplatin (Fig. 5A, B). *AKT3* encodes a member of the AKT serine/threonine protein kinase family. As a middle-stream regulator of IGF1 (insulin-like growth factor 1) signalling, AKT3 plays a key role in signalling and regulating cell survival in insulin signalling, angiogenesis and tumour formation. Cisplatin-induced damage to this gene could trigger a series of downstream changes, inducing apoptosis and death of cancer cells [31].

## DISCUSSION AND REMARKS

Cisplatin is a genetically toxic anticancer drug, but its gene targets or precise gene loci are largely unknown, limiting understanding of the cellular impact of DNA damage induced by the drug. In this work we have mapped genome-wide cisplatin-damage to gene loci, and further elucidated the associated core signalling pathways as well as diseases and toxicity by bioinformatics analysis using the IPA programme.

Two methods, termed “cisplatin-seq” [11] and “damage-seq” [12], have been previously developed to map the cisplatin-damaged sites in genomic DNA. Both these methods achieved single-base (nucleotide) resolution mapping of cisplatination sites on specific genomic regions such as promotors, transcription termination sequences and intergenic regions. However, the precise genes damaged by cisplatin and the biological consequences of the gene damage were not determined. Transcriptomics has been used in combination with the “damage-seq” method to map specific cisplatin-damaged gene loci across mouse organs [13]. Due to the tissue-specificity of transcriptomics, only the genes transcribed in specific tissues could be mapped for cisplatin lesions.

There are a number of chemical genetic strategies, in particular chemical proteomic approaches, that can uncover gene targets for cisplatin. These have produced significant advances in elucidating mechanisms of drug action. However, similar to the transcriptomics approach, they do not fully reveal cisplatin-damaged gene loci throughout the whole genome because they have focussed on discovering gene targets of cisplatin *via* measuring the difference between gene expression in specific types of cancer cells or tumour tissues upon cisplatin exposure. For example, Jimenez and co-workers found that UGGT1, COL6A1 and MAP4 can be used as potential biomarkers for determining cisplatin response, by proteomic analysis of non-small cell lung cancer cells [32]. Also Wilmes et al. have used integrated omics techniques to study differentially-expressed proteins and related pathways for cisplatin nephrotoxicity [28]. In this work, by integration of gene sequencing with bioinformatics analysis, our data provide three innovative findings. Firstly, cisplatin attacks 440 protein kinase genes with FEG >1.5 (Supplementary Table S5), covering 85% of consensual protein kinase genes in the human genome [18]. Secondly, cisplatin also induced damage on 110 protein phosphatase genes, accounting for 81% of the 136 reported protein phosphatases in the human genome (Supplementary Table S6). Last but not least, amongst the 14971 identified cisplatin-damaged protein-encoding genes, 418 genes are in the 628 prior gene targets for cancer treatment reported recently [20] (Supplementary Table S7).

Cisplatin-damaged genes reduced the expression of a series of protein kinase genes, bioinformatics analysis revealed that they are associated in a series of important signalling pathways such as protein kinase A (PKA) (Fig. 4A, Supplementary Table S9), as verified by Western Blot assays (Fig. 5). Protein kinases, accompanied by phosphatases, regulate protein phosphorylation, the most common and important protein post-translational modification, involved in many natural biochemical pathways and cellular processes [33]. Since abnormal protein phosphorylation is closely related to cancer development, protein kinases are major drug targets for cancer therapy [34]. A number of small molecule protein kinase inhibitors (PKIs), e.g. gefitinib, imatinib and dasatinib [35], have been developed and approved for clinical treatment of various cancers. Given our findings that protein kinase genes are highly susceptible to cisplatin attack, and that the cisplatin-damaged protein kinase genes are highly associated with 320 core signalling pathways, our work unambiguously demonstrates that cisplatin acts as a broad-spectrum protein kinase inhibitor by inducing lesions in protein kinase genes. The cisplatin-damaged protein phosphatase genes includes a series of protein tyrosine phosphatase receptor (PTPR), such as *PTPRD/F/M/E/J/U/S/T,* with FEG of > 50. This may explain why intracellular response to cisplatin activates rather than inhibits Ataxia Telangiectasia mutated (ATM)/ATM and Rad3-related (ATR)/DNA-protein kinase (PK) dependent pathways [36,37] while it induces damage on protein kinase genes as described above. Taken together, cisplatin damage on protein kinase and phosphatase genes is genome-wide, implying that cisplatin exerts its pharmacological functions by disturb the signalling of protein phosphorylation in cancer cells, with the potential to reprogram protein phosphorylation proteome-wide.

We also found that mitochondrial genes might not be the major targets for cisplatin, possibly because mitochondrial genes are usually short and have a low density of guanine, the major binding site of cisplatin on DNA. This result is consistent with the widely accepted knowledge that nuclear DNA (nDNA) is the major target of cisplatin [3,4]. However, some reports have suggested that mitochondrial DNA (mDNA) is more susceptible to cisplatin attack than nDNA [5,11]. In contrast, those studies used different cell types (HeLa ovarian cancer cells [11] and immortalized lymphocytes [12]), compared to A549 lung cancer cells used here, and different doses of cisplatin and times of treatment.

Importantly, we found significantly reduced expression of *SPAG9* by cisplatin-induced lesions, notably contributing to cisplatin-induced apoptosis as verified by siRNA silencing (Fig. 5C – F). As shown in Supplementary Fig. S22A, *SPAG9* (JIP-4) activates JUN/MAP kinase signalling [27,38], and is associated with various types of cancers, e.g. anaplastic thyroid carcinoma [39], hairy cell leukaemia [40] and intestinal gastric adenocarcinoma [41]. It also plays a role in several important biological processes, such as fertility [42], differentiation of skeletal muscle cells [43], and differentiation of neurons [44]. Moreover, the *SPAG9* gene is regulated by ligand-dependent nuclear receptor ESR1 [45], transmembrane receptor CAV1 [46], transcription factor ZNE217 [47] and transporter BSG [48], and regulates mainly ERK, JNK and p38 MAPK signalling [49] (Supplementary Fig. S22B). JIP-4 functions as a scaffold protein that structurally organizes MAPKs and mediates c-Jun-terminal kinase signalling. It has been demonstrated that *SPAG9* is associated with tumour growth, migration, and invasion in renal cell carcinoma [50]. Sinha et al. discovered that down regulation of *SPAG9* by siRNA silencing reduces growth and invasion potential of triple-negative breast cancer cells [51]. Because *SPAG9* specifically expresses the upstream mediator JIP-4 of p38 MAPK signalling in high abundance in testis [52,53], the cisplatin damage on *SPAG9* may accounts for its high cure rate for testicular cancer, which deserves further investigation.

Cisplatin is the most successful metallo-anticancer drug in clinic. Over 50% of the chemotherapy treatments use cisplatin related compounds. In contrast to many organic drugs, metallo-drugs are usually multi-targeted, and it is essential to understand the nature of their target sites to elucidate their systems pharmacology [54]. It is evident from our work that cisplatin targets a very wide variety of kinase and phosphatase genes. This provides novel insights into the mechanism of action of this drug and a basis for design of next generation of platinum anticancer drugs. The design would involve drug delivery systems that target specific protein kinase genes which play crucial roles in the development of particular cancers [55], and can involve minimizing non-specific attack on protein kinase genes of healthy cells [2,16].

## DATA AVAILABILITY

The raw sequencing data are deposited on Mendeley Data, and available on https://data.mendeley.com/datasets/k9sm56gj5y/1, https://data.mendeley.com/datasets/2tv7ntcgh6/1, https://data.mendeley.com/datasets/4zj3kz34c5/1, https://data.mendeley.com/datasets/dmyfkn7pnz/1, https://data.mendeley.com/datasets/dw3v4sm9nb/1, https://data.mendeley.com/datasets/xb6s7vzcwm/1, https://data.mendeley.com/datasets/75dyzmvbrz/1, https://data.mendeley.com/datasets/s7jbxvwf4z/1, and https://data.mendeley.com/datasets/fv6f9rpg5m/1.

The flow cytometry data contained in this article are deposited on FlowRespository with a Repository ID of FR-FCM-Z45H on https://flowrepository.org/experiments/4273.

## Abbreviations

A549: non-small cell lung cancer cell line
AKT: RAC-gamma serine/threonine-protein kinase
CDGs: Cisplatin-damaged genes
CS: control sample
EGFR: epidermal growth factor receptor
FEG: fold-enrichment of a gene
FEP: fold-enrichment of a peak
GAPDH: Glyceraldehyde-3-phosphate dehydrogenase
GSH: glutathione
HMGB1: human high mobility group box 1
HMGB1a: HMGB1 box a
ICP-MS: Inductively Coupled Plasma-Mass Spectrometry
IGF1: insulin-like growth factor 1
IPA: Ingenuity Pathway Analysis
JIP-4: C-jun-amino-terminal kinase-interacting protein 4
MAPK: mitogen-activated protein kinase
mDNA: mitochondrial DNA
MERTK: c-mer proto-oncogene tyrosine kinase
NCCIT: human testicular cancer cell line
nDNA: nuclear DNA
NFAT: nuclear factor of activated T cells
NGS: Next Generation Sequencing
NGS: Next Generation Sequencing
PI3K: phosphatidylinositol 3’-kinase
PK: protein kinase
PKA: protein kinase A
PKI: protein kinase inhibitors
PS: positive sample
PTPR: protein tyrosine phosphatase receptor
SP: sperm motility
SPAG9: sperm associated antigen 9
XR-seq: Repair-seq

## Author contributions

F.Y.W. and Y.Zhao. conceived the project. L.Y.Q., Y.Z.H, Q.L., and Y.X. carried out the expression of HMGB1a and construction of HMGB1a-functionilizaed microprobes. L.Y.Q., B.B.X., Y.L.H., Y.Y.Z., X.J.L., P.J.S., Y.Zhao. and F.Y.W. analysed sequencing data. L.Y.Q., W.C.Y. and X.K.L. performed bioinformatics analysis and Western Blot assays. L.Y.Q., F.Y.W. P.J.S., Y.Zhao. and S.J.W. wrote and revised the manuscript with inputs from all authors.

## Additional information

**Supplementary information** is available for this paper at …….

**Correspondence and requests for materials** should be addressed to F.Y.W.,Y.Z., P.J.S. or S.J.W

