## Supplementary information for "Genomic Mapping Reveals Cisplatin Disruption of Protein Phosphorylation Signalling Genome-Wide"

**Table of Contents**

**Experimental Section**

**Platination level and the quality check for the DNA fragments**

**Supplemental Figures S1 – S23**

**Experimental Section**

**Materials and Methods**

**Culture of *Escherichia coil***

*E. coli* strain BL21 with HMGB1a-encoding plasmid (DE3) (Tian Gen Biotech, China) was cultivated in Luria-Bertani (LB) medium with 50 μg/mL kanamycin. A single colony picked from a fresh plate was used to inoculate 2YT medium (containing Bacto^TM^ Peptone (16 mg/mL), Bacto^TM^ Yeast Extract (10 mg/mL, Becton, Dickinson and company, USA), sodium chloride (5 mg/mL, Beijing Chemical Works, China) and kanamycin (50 μg/mL, Solarbio, China)) overnight at 37 °C and 180 rpm. Then the cells were transferred to a large volume of fresh 2YT medium at 37 °C and 180 rpm. When the OD_600_ value of the cells reached 0.5 – 1.0, IPTG (Sigma-Aldrich, USA) was added at a final concentration of 1 mM to induce gene expression. After incubation at 37 °C and 180 rpm for further 12 h, the cells were harvested and washed once with PBS (Solarbio, China), and frozen at −80°C for at least one hour to improve sonication.

**Culture of cancer cell lines**

The A549 human non-small cell lung cancer cell line (National Infrastructure of Cell Line Resource, Beijing, China) and NCCIT testicular cancer cell line (Fukesai Biological Technology Co., Ltd, Nanjing, China) were individually cultured in DMEM medium (Gibco) supplemented with 10% FBS (Gibco) and 1% penicillin and streptomycin (GE Healthcare) at 37 °C. When the cell density reached confluency, the culture medium was replaced by fresh DMEM in the absence or presence of cisplatin at various concentrations (50 μM for A549, 12 μM for NCCIT) for 24 h. Then the cells were harvested and washed twice by PBS. Whole cell protein or genomic DNA were extracted from different cells with different extraction kits.

**Expression and purification of HMGB1a**

The cDNA of human HMGB1 domain A (HMGB1a, Supplementary Fig. S23A) was synthesized by Proteintech (Rosemont, USA) and cloned into pET-28a plasmid (Supplementary Fig. S23B). The resulting plasmid encoding a his_6_-tag was transfected into BL21 (DE3) *E. coli* cells (Tian Gen Biotech, China). The cells were grown in a small volume of 2YT medium containing Bacto^TM^ Peptone (16 mg/mL), Bacto^TM^ Yeast Extract (10 mg/mL, Becton, Dickinson and company, USA), sodium chloride (5 mg/mL, Beijing Chemical Works, China) and kanamycin (50 μg/mL, Solarbio, China) overnight at 37 °C and 180 rpm. Then the cells were transferred to a large volume of fresh 2YT medium at 37 °C and 180 rpm. When the OD_600_ value of the cells reached 0.5 – 1.0, IPTG (Sigma-Aldrich, USA) was added at a final concentration of 1 mM to induce gene expression. After incubation at 37 °C and 180 rpm for further 12 h, the cells were harvested and washed once with PBS (Solarbio, China), and frozen at −80°C for at least one hour for improved sonication. The deep frozen cells were resuspended in 10 mL cold lysis buffer (PBS and 5 mM imidazole, pH 7.4), and lysed by sonication for 20 min at 0 °C, followed by centrifugation for 30 min at 4 °C and 12000 rpm. The supernatant was purified by His Trap^TM^ FF crude column (GE Healthcare) which was eluted first by 50 mM imidazole (Innochem), then by 300 mM imidazole in phosphate buffer. The eluted HMGB1a containing a his_6_-tag was characterized by SDS-PAGE (Supplementary Fig. S23C) and mass spectrometry (Supplementary Fig. S23D) .

**Assembly and characterization of HMGB1a-functionalized affinity microprobes**

To construct affinity microprobes for cisplatin crosslinked DNA, 2.1 mg purified HMGB1a was incubated with 7.5 mg nickel-modified magnetic beads (MAg25K/IDA Ni, Enriching Biotechnology, Shanghai, China) suspended in 1 mL PBS, and rotated for 1 h at 4 °C. The beads were then separated by magnetic force and washed twice with PBS for 10 min. As shown in Supplementary Fig. S23E, the absorbance of the supernatant over the wavelength range 250 – 300 nm significantly decreased due to loading of HMGB1a onto the magnetic beads. This indicated successful functionalization of the magnetic beads by HMGB1a.

**Cell culture and genomic DNA extraction**

The A549 human non-small cell lung cancer cells obtained from the Centre for Cell Resource of Peking Union Medical College Hospital were cultured in DMEM medium (Gibco) supplemented with 10% FBS (Gibco) and 1% penicillin and streptomycin (GE Healthcare) at 37 °C. When the cell density reached confluency, the culture medium was replaced by fresh DMEM containing 50 μM cisplatin (Beijing Ouhe Technology, China), and the cells were incubated at 37 °C for 24 h. Then the cells (~10^7^) were harvested, and genomic DNA was extracted by whole genome extraction kit (Tian Gen Biotech). The DNA extracts were crushed into fragments between 250 to 750 bp with an ultrasonic breaker (Scientz-IID ultrasonic homogenizer, SCIENTZ, Ningbo, China). At the same time, a similar amount of cells without exposure to cisplatin were harvested for genomic DNA extract and fragmentation. The DNA fragments from the cells treated with cisplatin were considered as a positive sample and those from the cells without cisplatin treatment as a control sample for capture and sequencing of cisplatin-damaged genes. An inductively coupled plasma-mass spectrometer (ICP-MS, Agilent 7700) was used to determine the Pt concentrations of the extracted DNA obtained from positive and control samples.

**Capture of cisplatin-bound DNA fragments**

The same amounts (74 μg) of DNA fragments gained from cells treated and untreated with cisplatin were individually re-dissolved in 10 mM Tris-HCl (pH 7.4, Beijing Biodee Biotechnology), and added into the HMGB1a-functionlized magnetic beads suspended in PBS, followed by rotation for 1 h at room temperature. The beads were then separated by magnetic force and washed twice with 1 mL 10 mM Tris-HCl. The captured HMGB1a-DNA complexes were eluted with 1 mL of buffer containing 20 mM sodium phosphate, 500 mM NaCl and 400 mM imidazole (pH 7.4) *via* rotation at room temperature for 40 min. As shown in Supplementary Fig. S23F, G, the absorbance of the supernatant over the wavelength range 250 – 300 nm significantly decreased subject to incubation with HMGB1a-functionlized magnetic beads, providing evidence for the capture of DNA fragments by the HMGB1a-based affinity microprobes.

Each eluted HMGB1a-DNA complex sample was centrifuged and lyophilized, and then re-dissolved in 500 μL deionized water, followed by addition of 500 μL mixture of phenol (Solarbio) – chloroform (Beijing Chemical Works) – isopropanol (Concord Technology, Tianjin) (25: 24: 1). The resulting mixture was shaken firstly for 10 s, and kept at room temperature for 2 min, then centrifuged at 12000 rpm and 4 °C for 7 min. The supernatant was mixed with 500 μL of a mixture of chloroform-isopropanol (24: 1), shaken again for 10 s, kept at room temperature for 2 min, and then centrifuged at 12000 rpm and 4 °C for 7 min. The isolated supernatant was mixed with 1500 μL pre-chilled absolute ethanol (Concord Technology, Tianjin) and 50 μL sodium acetate solution (3 M, Beijing Chemical Works), kept at −20 °C overnight, and then centrifuged at 12000 rpm and 4 °C for 7 min. The supernatant was carefully removed, and the residues were left at room temperature for drying. Finally, the purified DNA fragments were re-dissolved in an appropriate amount of TE solution (Solarbio) for sequencing.

**Sequencing and gene-mapping**

The next-generation-sequencing (NGS) of the captured and purified DNA fragments described above was performed by Shanghai Sangon Biotech Ltd. The DNA fragments were firstly characterized by 1% agarose gel electrophoresis for quality check, and then fragmented to about 500 bp by S220 Focused-ultrasonicator and recovered for DNA library construction. Library construction was performed following the instruction of NEB Next® Ultra^TM^ DNA Library Prep Kit for Illumina^®^ (<https://international.neb.com/products/e7370-nebnext-ultra-dna-library-prep-kit-for-illumina#Protocols,%20Manuals%20&%20Usage>). Since cisplatin binding to DNA fragments inhibits DNA replication and synthesis[1-3], the PCR expansion time was prolonged to 30 min. The constructed DNA libraries were characterized by a 2% agarose gel, and then each positive or control sample was sequenced by Illumina®, providing at least 20 G of data.

The raw Illumina sequences were filtered for quality control by using Fastp (0.22.0) (fastp -q 15 -u 40 -n 5 -e 0 -l 35), and the clean reads were aligned to the reference human genome (hg19; National Centre for Biotechnology Information [NCBI]) by using the Burrows-Wheeler Aligner (BWA) software program (0.7.17-r1188) with default parameter [bwa mem -M <genome> <R1> <R2>] to acquire SAM files. Then, Samtools (1.6) was used to sort out the SAM files to BAM files in which only uniquely mapped nonduplicate reads were retained. High resolution ChIP-Seq peaks predictions were conducted by MACS2(2.1.2) program with a p-value cutoff of 0.001 and Mfold values adjusted to build the model, [macs2 callpeak -c <treat BAM > -t <control BAM> -q 0.05 -f BAMPE -g hs]. Finally, the R-Chipseeker package was employed to annotate and visualize all peaks identified.

To calculate the fold-enrichment of a cisplatin-damaged gene (CDG), we first calculated the fold-enrichment of all peaks mapped to the gene by using equation 1.

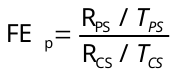
 (1)

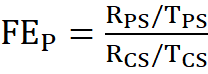

Where R_PS_ and R_CS_ are read (tag) counts for a peak detected in a pair of positive sample (PS) and control sample (CS), respectively, and T_PS_ and T_CS_ are total read (tag) counts of all peaks detected in the pair of positive sample (PS) and control sample (CS), respectively. A peak was considered unreliable and then abandoned if it matched multiple genes. A peak with an fold-enrichment (FE_p_) <1.5 was also taken as unreliable and discarded[4]. Thus, the fold-enrichment of a CDG is the sum of fold-enrichment of all peaks mapped to the gene, equation 2.

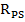

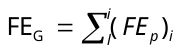
 (2)

Where i represents the number of peaks mapped for a gene in a positive sample/control sample pair.

Taking into account that cisplatin kills cells by attack on DNA, it might be expected that more cisplatin-damaged genes would be detected in the dead cells of positive samples. However, our sequencing data showed that there was no significant difference between the total numbers and fold-enrichments of cisplatin-damaged genes mapped from the DNA fragments collected from living cells alone and from living cells and dead cells together after 24 h of incubation with 50 μM cisplatin (data not shown). The other software and databases used for gene screening and localization are listed in Supplementary Table S17.

It is notable that due to the large difference in the length of genes, the FE_G_ of a CDG based on the fold-enrichment of all peaks to the gene does not correspond simply to the base coverage of all identified peaks for the gene (Supplementary Table S2). For example, 198 peaks were identified for gene named *TRAPPC9,* which has the highest FE_G_ (476.4) but with a low base coverage of 15.5%. In contrast, for microRNA gene *MIR182* with a FE_G_ of 7, only 3 peaks were identified, but the base coverage of these peaks to the whole gene of *MIR182* was 74.7%. With regard to this, the FE_G_ in this work is an indicator only of CDG-mapping confidence, and not of the degree of cisplatin damage to a gene.

**Bioinformatics Analysis**

Ingenuity Pathway Analysis (IPA, QIAGEN, 2025)[5] was used to analyse the core pathways, diseases, toxicities and functions associated with the cisplatin damage. IPA analysis maps each gene to the corresponding molecule in the Ingenuity Pathway Knowledge Base, which is available at the Ingenuity System’s web site (http://www.ingenuity.com).

**CCK-8 assay**

The survival rates of *SPAG9*-selicenced and cisplatin-treated NCCIT cells were measured using the CCK-8 (Cell Counting Kit-8) assay (MedChemExpress, USA). Briefly, the NCCIT cells (1×10^3^ cells/per wells) were plated at a density of 2000 cells per well in 100 μL of media into 96-well plates. After 24 h for adherence, the cells were exposed to 12 μM cisplatin for 24 h or transfected with the siRNA against *SPAG9* and grown for 48 h. Then, 10 μL CCK-8 was added into the medium and the cells were grown for further 4 h. Finally, the optical density (OD) for each well was measured using a microplate reader (SpectraMax M5 Molecular Devices Corporation) at the wavelength of 450 nm. The survival rate (SR, %) of cells was calculated based on equation (1):

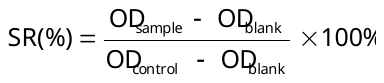
 (3)

All reported values of SR are averages of three independent experiments and expressed as mean ± SD (standard deviation).

**Flow cytometric analysis**

The apoptosis rate of NCCIT cells subject to cisplatin treatment and *SPAG9* silencing was determined by Annexin V / propidium iodide (PI) double-staining assay following the Apoptosis Detection Kit I protocols (BD Pharmingen, USA). The NCCIT cells (1×10^3^ cells/well) were plated into a six-well plate. Twenty-four hours later, NCCIT cells were transfected with siRNA against *SPAG9* for 48 h. Another group of cells was treated with 12 μM cisplatin for 24 h. Then, the supernatant was removed, and cells were transferred to tubes and centrifuged at 1000 rpm for 3 min. Thereafter, the cells were re-suspended in 1× Annexin V binding buffer, and incubated with 5 μL Annexin-V and 5 μL PI conjugate for 15 min in the dark prior to FACS analysis. The FACS assays were performed on a Calibur flow cytometer (BD，Franklin Lakes, New Jersey, US), of which the FL1 channel was used to record the intensity of annexin V-FITC staining and FL2 channel to record the intensity of PI staining. The data were quantified by Sell Quest software (BD, Franklin Lakes, New Jersey, US).

**Quantification of protein and DNA concentration**

The concentrations of purified protein HMGB1a and extracted genomic DNA were determined by NanoDrop OneC (Thermo Fisher Scientific Inc.). The concentrations of whole cell proteins from different cells were determined using a BCA kit (Beyotime Biotech Inc.).

**Western Blot assays**

The A549 human non-small cell lung cancer cell line (National Infrastructure of Cell Line Resource, Beijing, China) and NCCIT testicular cancer cell line (Fukesai Biological Technology Co., Ltd, Nanjing, China) were individually grown in the presence and absence of cisplatin at various concentrations (50 μM for A549 and 12 μM for NCCIT) for 24 h. Different doses of cisplatin were applied to treat the cell lines because of the different sensitivity of the cells to cisplatin, the IC_50_ value of 5.8 μM (24 h) for NCCIT cell line and 10.1 μM (48 h) for A549 cell line[6]. The cells were then harvested for extracting whole cell proteins using a total protein extraction kit (BestBio, China). The concentration of each protein extract was determined by the standard BCA assay (Beyotime, China). To determine the change in expression level of a specific protein due to gene damage induced by cisplatin, the same amount of whole cell proteins extracted from a positive sample and a control sample were individually boiled with the gel-loading buffer (4×LDS, Genscript, Nanjing, China), and loaded onto a 4 – 12% gradient SDS-PAGE gel (Genscript) for electrophoresis. The separated proteins were transferred to a PVDF membrane (Millipore, 0.2 μm). The membrane was blocked in 5% defatted milk powder (Biofroxx) dissolved in 0.1% TBST, of which 1 L contained 2.4 g Tris base (Innochem), 8.8 g NaCl and 0.1% (v/v) Tween 20 (Beijing Topbio Science & Technology), at 4 °C overnight, and then incubated with respective primary antibodies (Abcam) in appropriate dilutions at room temperature for 2 h. Thereafter, the membranes were washed three times with 5% nonfat dry milk in 0.1% TBST for 30 min each time and incubated with horseradish peroxidase-conjugated secondary antibody (Abcam) at room temperature for 1 h. After washing twice with 5% defatted milk powder in 0.1% TBST and once with 0.1% TBST, the protein bands were visualized by enhanced chemiluminescence (Beyotime, China), and the optical densities were determined using an image analyser (Image J).

For siRNA silencing of *SPAG9* experiments, the NCCIT testicular cancer cell line were transfected with siRNA against *SPAG9* (sequence 5’-3’: rGrGrArGrCrArGrArUrUrUrArCrUrArGrGrArATT, Suzhou Biosyn Biotechnology Co., Ltd.), or not, for 48 h. The cells were then harvested for extracting whole cell proteins using a total protein extraction kit (BestBio, China). The concentration of each protein extract was determined by standard BCA assay (Beyotime, China). To determine the change in expression level of JIP-4 due to gene silencing by siRNA, the same amount of whole cell proteins extracted from a positive sample and a control sample were individually analysed by using the same method for Western Blot assays described above.

**Calculation of fold-change of protein expression**

We carried out three technical repetitions of Western Blot assays. The optical densities of protein bands were determined using an image analyser (Image J). The fold-change (FC) of a protein was calculated using equation 4.

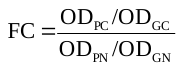
 (4)

Where OD_PC_ and OD_GC_ are optical densities of a protein band and the GAPDH (the reference) band in a sample treated with cisplatin, and OD_PN_ and OD_GN_ are optical densities of the protein band and the GAPDH band in a sample without cisplatin treatment, respectively.

**Statistical analysis**

All data were analyzed with Origin 8.0 and Microsoft Offices Excel 2021 software. Statistically significant differences were determined using unpaired t-tests, Student′s t-tests, Fisher's exact test right-tailed, as applicable and according to test requirements. A two-sided *p* value < 0.05 was considered statistically significant. * indicates *p* < 0.05, ** *p* < 0.01, *** *p* < 0.001. No statistical methods were used to predetermine sample size. The experiments were not randomized and the investigators were not blinded to allocation during experiments and outcome assessment.

**Platination level and the quality check for the DNA fragments**

**Determination of the platination level of DNA fragments**

Inductively coupled plasma-mass spectrometry (ICP-MS) analysis for Pt showed that the platination level of genomic DNA extracted from cisplatin-treated cells reached 320 Pt atoms per 10^6^ base pairs (Fig. 2A), consistent with our previous report[7].

**Quality check for the cisplatin-bound DNA fragments**

Gel-electrophoresis data demonstrated that the size of the DNA fragments gained from both the PS and the CS ranged from 250 to 1000 bp, but the number of larger DNA fragments from positive samples was less than those from control samples (Fig. 2B). This is likely ascribed to platination, which could render genomic DNA more susceptible to sonication and more prone to aggregation[8]. Given that HMGB1a binds to both non-platinated and platinated DNA, it is not surprising that the same amount of the microprobes captured similar amounts of DNA fragments from positive and control samples (Supplementary Table S1). However, the subsequent sequencing revealed that there were significant differences between the DNA fragments enriched from the positive and control samples.

We constructed DNA libraries based on the captured DNA fragments. Cisplatin damage on DNA may stall DNA replication and/or cause base mismatch[3,9]. Therefore we attempted to remove Pt bound to DNA bases by using excess of glutathione (GSH) since Pt(II) has a high affinity for S-donor ligands[10]. However, we did not observe a significant difference between the amplification rates of GSH-treated and intact platinated DNA (data not shown). Previous reports have demonstrated that the replication of DNA can still reach a similar level for cisplatin-damaged DNA compared to intact DNA on extension of the replication time[3]. Therefore, we extended the PCR expansion time of both positive and negative samples to 30 min. The results suggest that the DNA sequences in DNA libraries of the positive samples and control samples have similar sizes (Fig. 2C), and that possible Pt-modification-induced bias for PCR amplification of DNA was circumvented (details in Materials and Methods).

**Supplemental Figures S1 – S23**

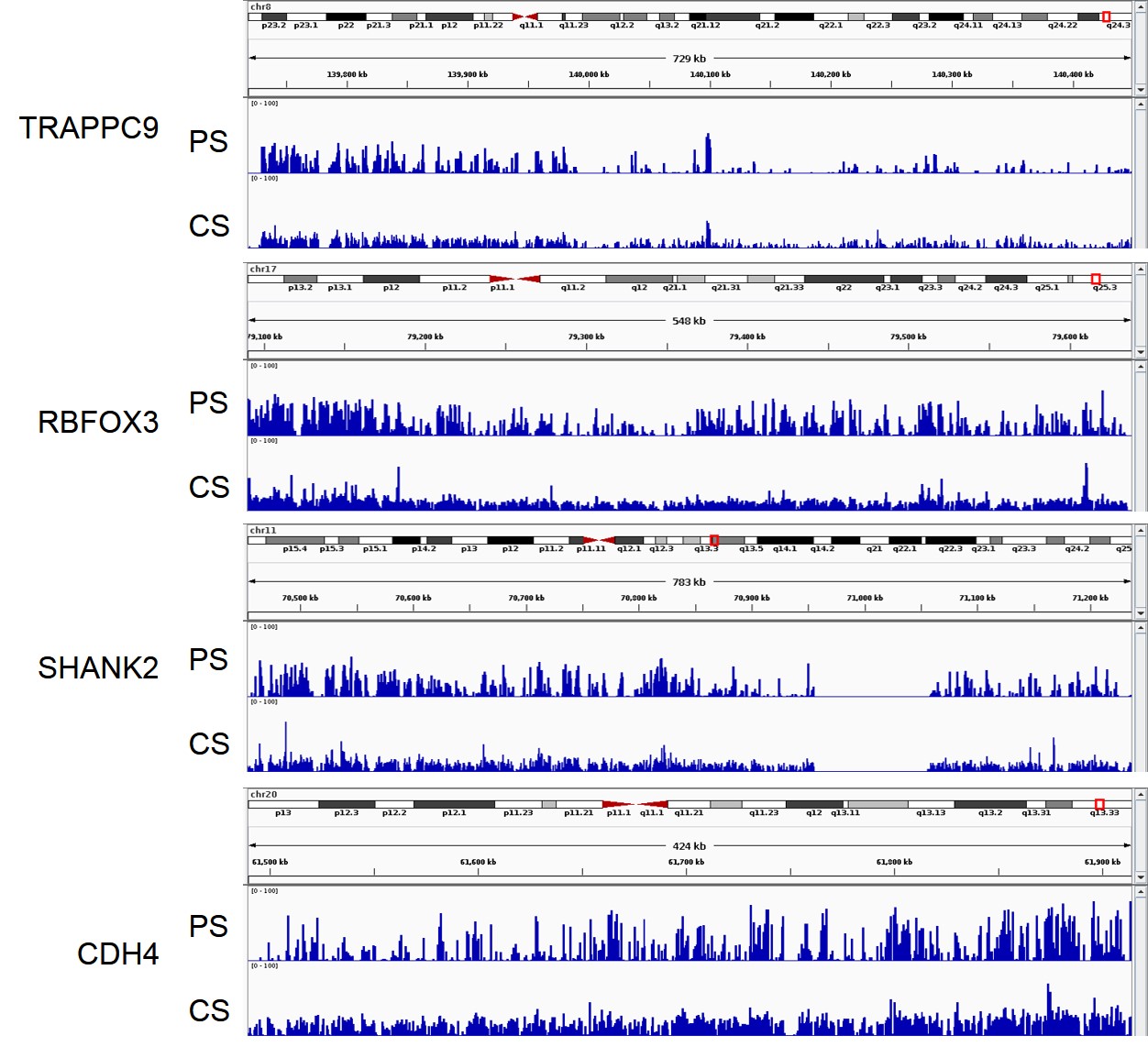

Supplementary Fig. S1. Reads for each peak mapped to four CDGs with the highest fold-enrichments in comparison with control. The x axis represents the location of a peak in the linear sequence of each gene, and the y axis represents the normalized read counts of the peak in the gene. PS: positive sample; CS control sample.

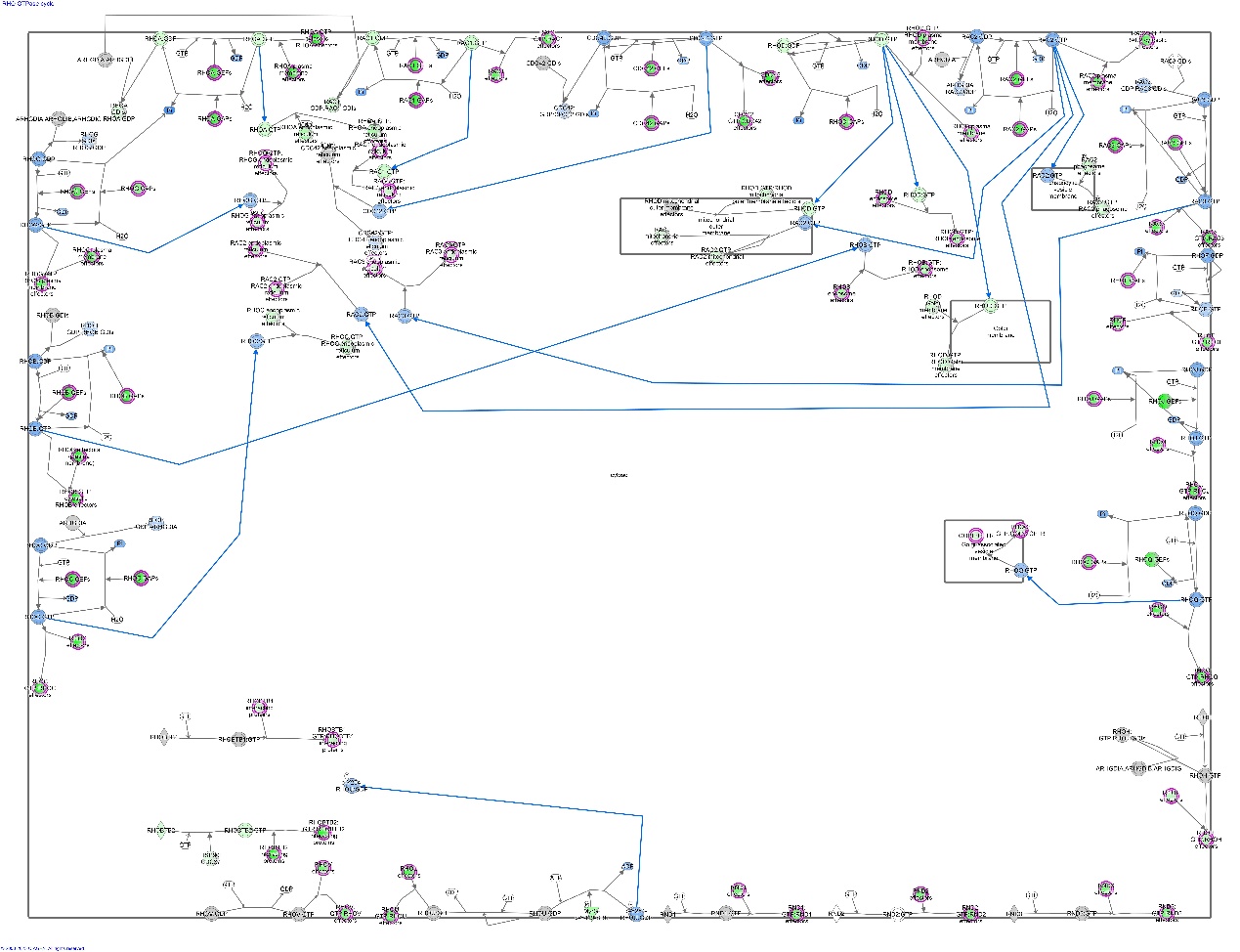

Supplementary Fig. S2. RHO GTPase cycle signalling pathway.

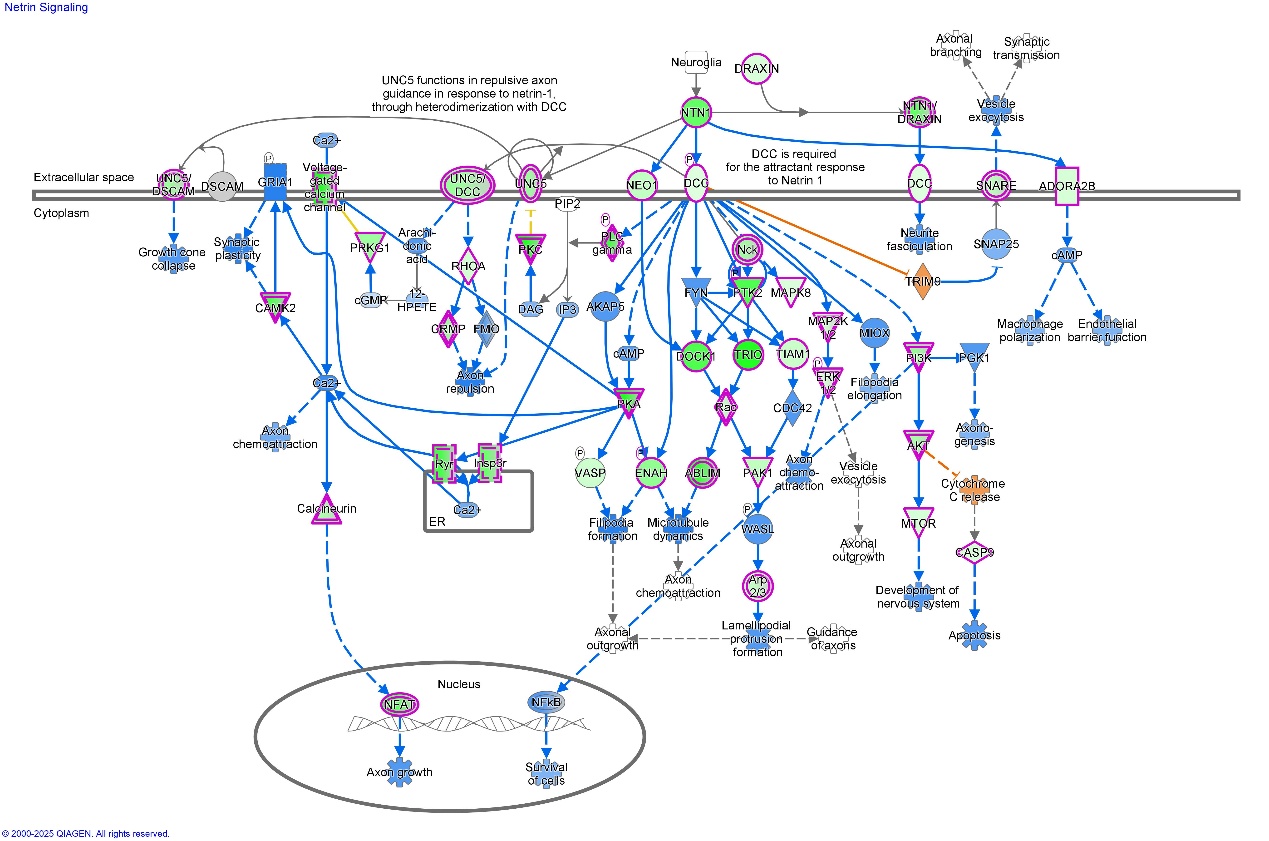

Supplementary Fig. S3. netrin signalling pathway.

A number of signalling pathways in the neural system are also associated with the CDGs, e.g. netrin signalling (*z*-score −8.1), synaptogenesis signalling pathway (*z*-score −10.8), myelination signalling pathway (*z*-score −8.7), glutaminergic receptor signalling pathway (*z*-score −10.3) (Fig. 4A, Supplementary Table S9 and Supplementary Figs. S3, S4, S6, S7). The high negative *z*-scores suggest that these signalling pathways are significantly inhibited by cisplatin damage on genes. This finding will stimulate exploration of whether and how cisplatin damage to neural signalling-related genes is related to its neurotoxicity[11,12].

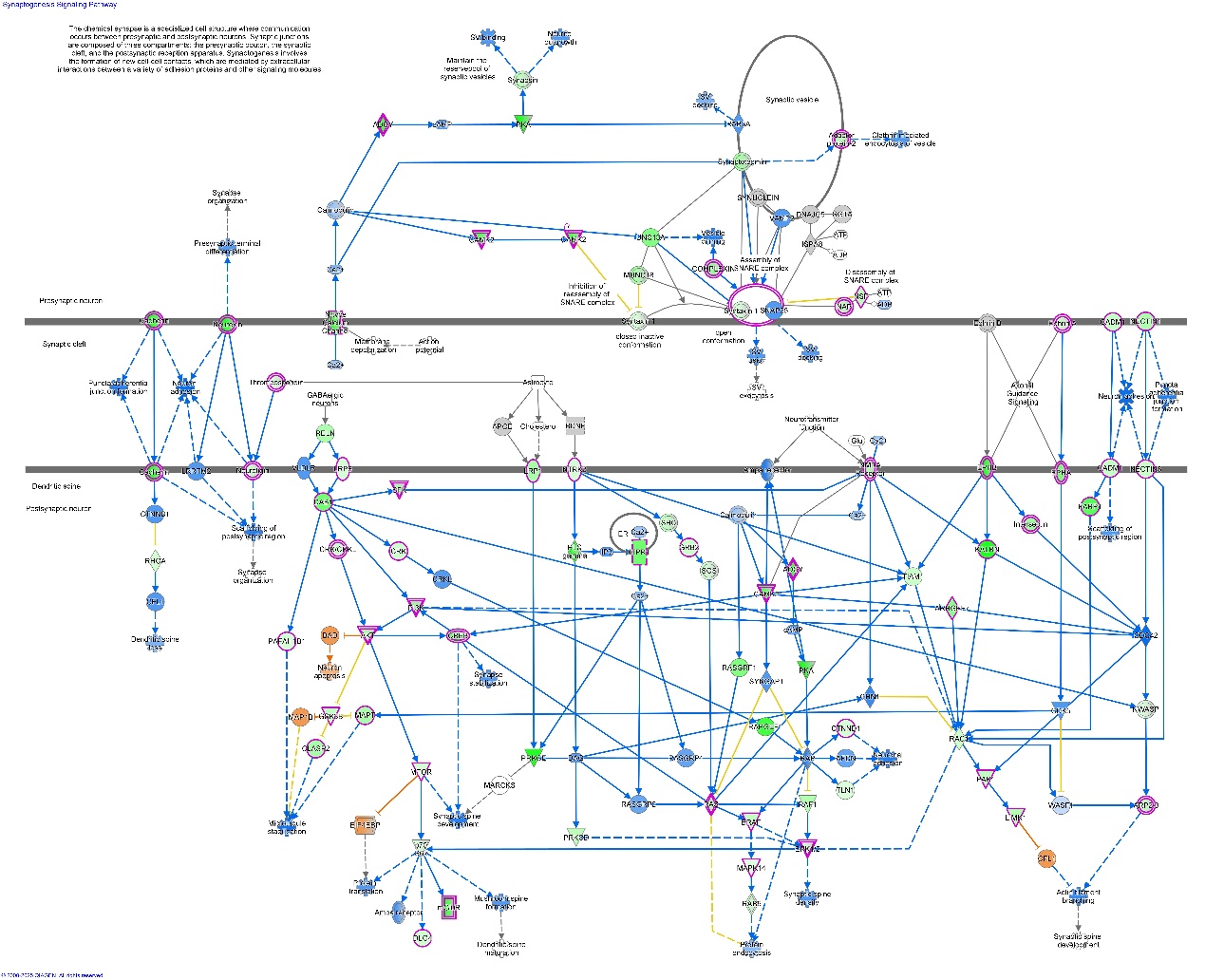

Supplementary Fig. S4. Synaptogenesis signalling pathway.

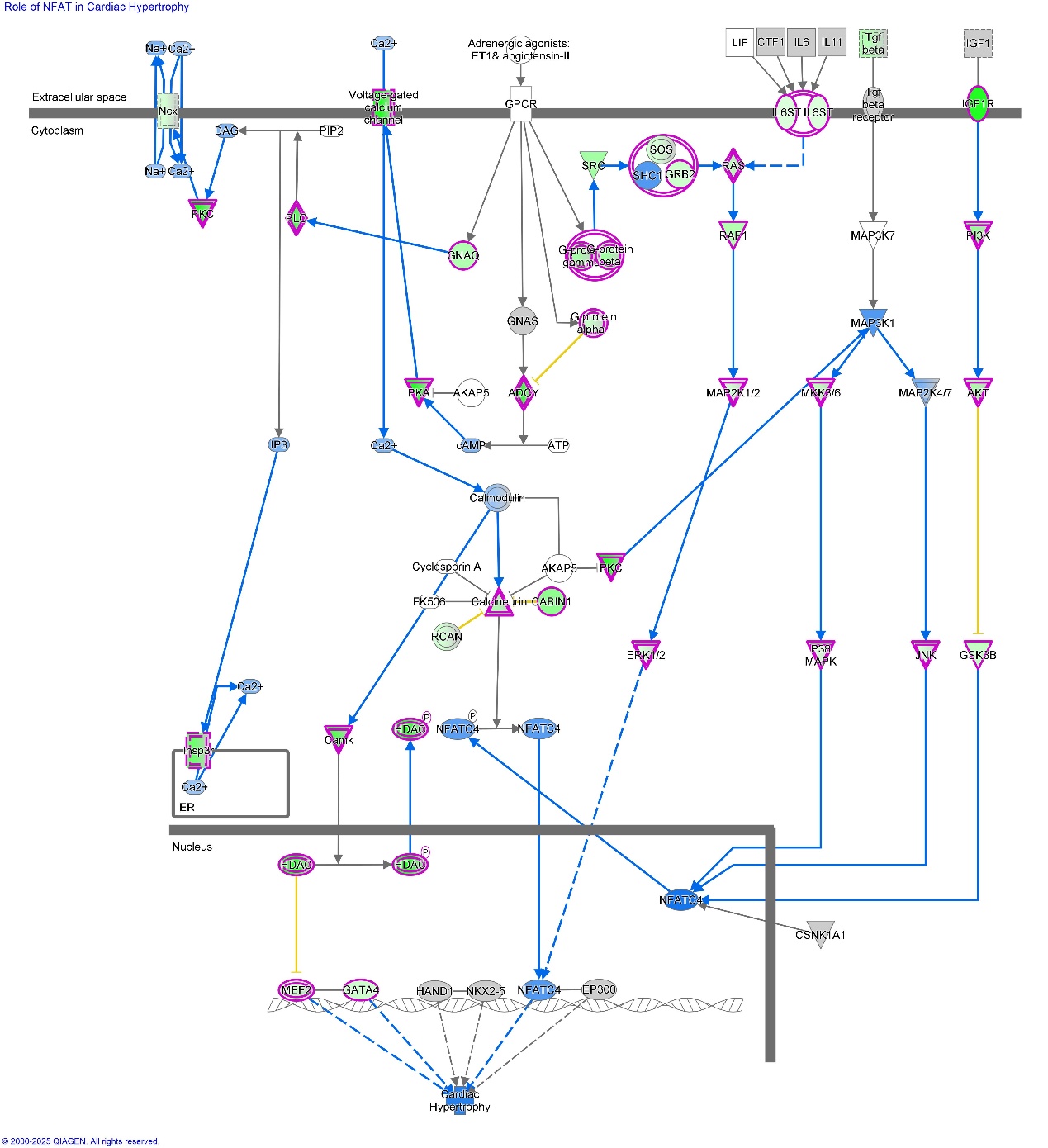

Supplementary Fig. S5. Role of NFAT in cardiac hypertrophy signalling pathway.

A series of cardiac related signalling pathways were also revealed. For example, IPA revealed that cisplatin-damaged genes are highly related to the role of NFAT in the cardiac hypertrophy and cardiac hypertrophy (enhanced) signalling pathway with −log *p* = 21.7 and 20.0, respectively (Supplementary Table S9 and Supplementary Fig. S5, S8). NFAT is the nuclear factor of activated T cells, as a transcription factor plays an important role in cardiac hypertrophic signalling which is mediated by the calcium-activated phosphatase calcineurin[13]. Calcineurin-NFAT signalling is linked to calcium signalling, with which 99 CDGs are highly associated and −log *p* = 18.0 (Supplementary Fig. S15), covering 50% of all the 198 genes involved in this signalling pathway (Supplementary Table S9). The calcium signalling pathway is predicted to be significantly inhibited by cisplatin damage to genes with a *z*-score of −7.25. The calcium channel genes *CACNA1A/B/C* and *CACNA2D3* are all damaged by cisplatin with FE_G_s of 213, 124, 133 and 112, respectively (Supplementary Table S2), suggesting that damage to these genes by cisplatin may be involved in cardiac hypertrophy, one of the clinical side effects of cisplatin.

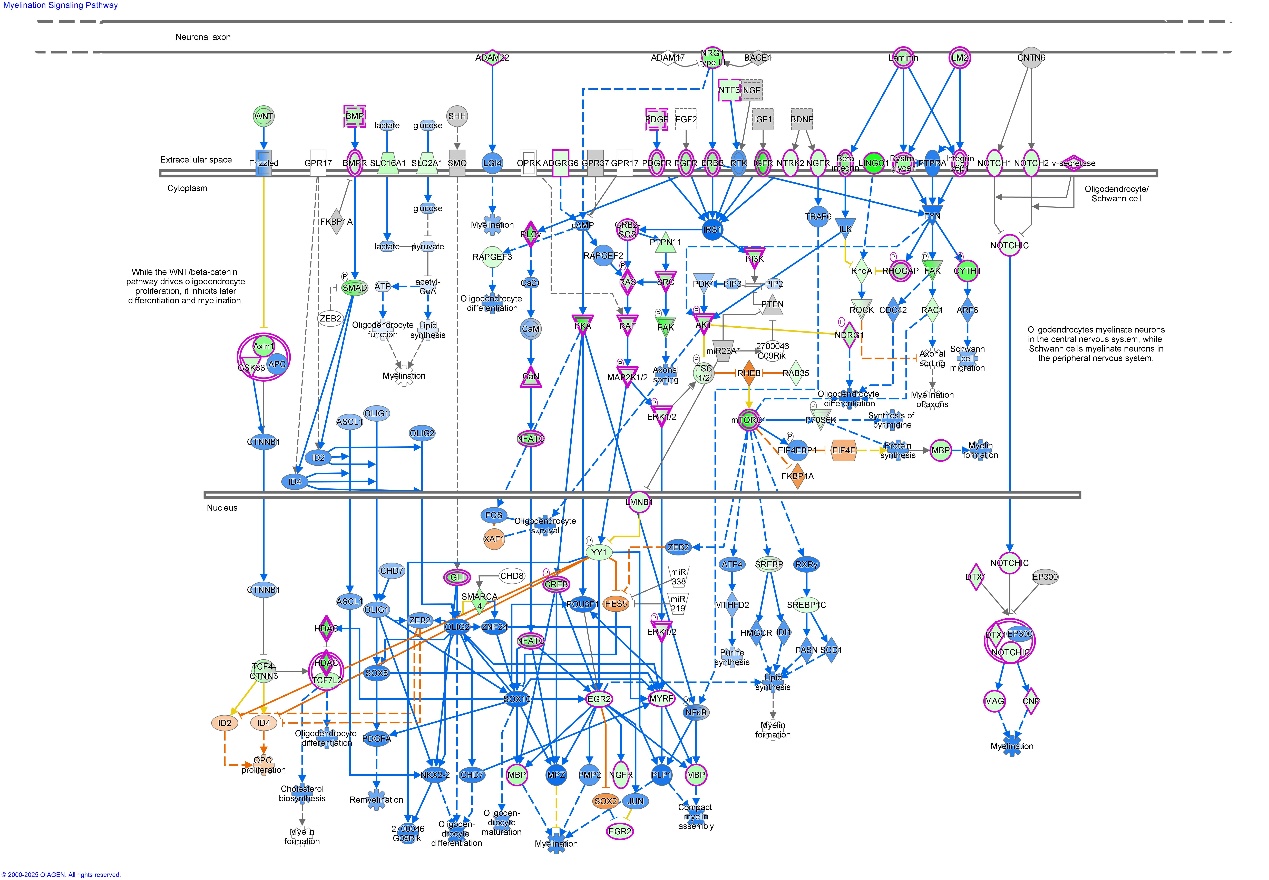

Supplementary Fig. S6. Myelination signalling pathways.

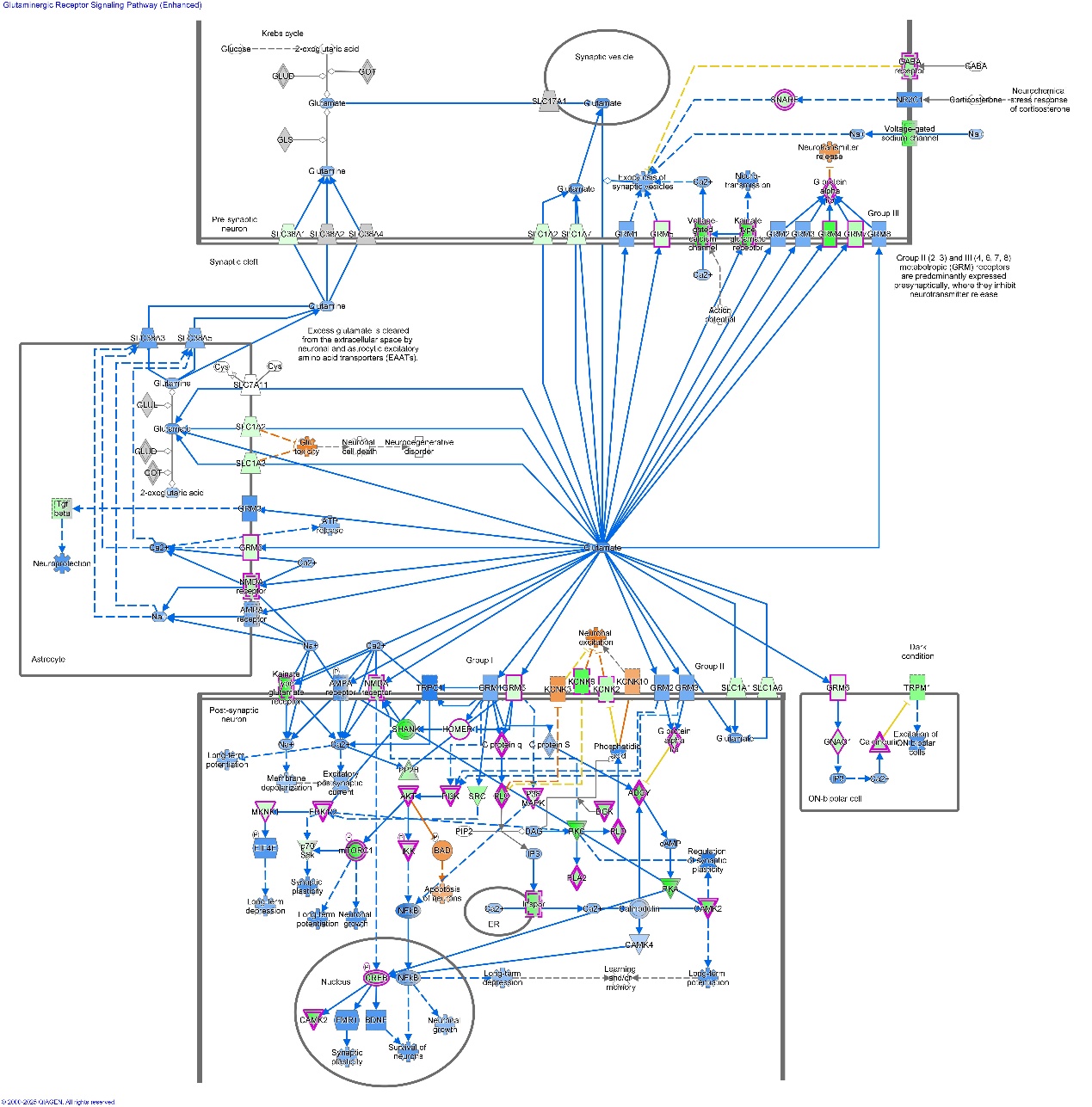

Supplementary Fig. S7. Glutaminergic Receptor Signalling Pathway.

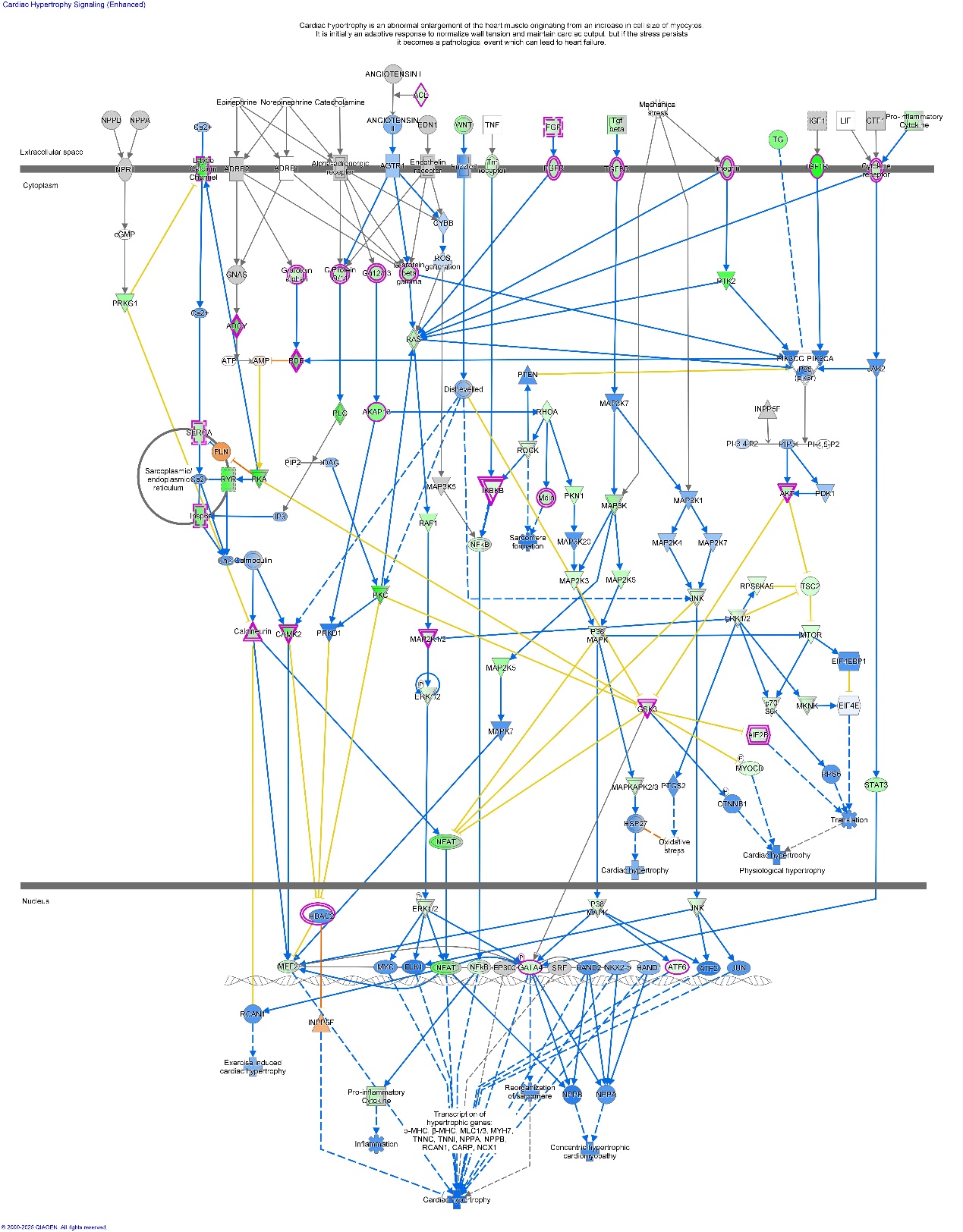

Supplementary Fig. S8. Cardiac Hypertrophy (Enhanced) Signalling Pathway.

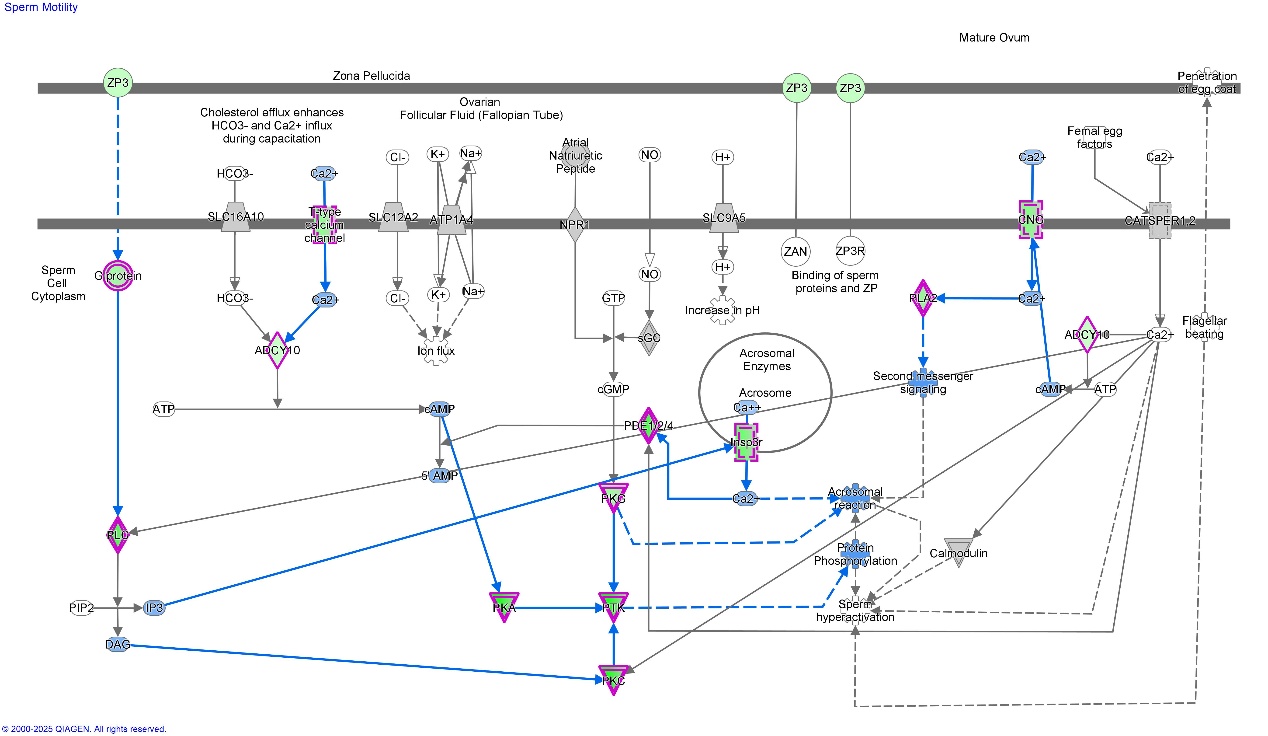

Supplementary Fig. S9. Sperm Motility Signalling Pathway.

One hundred and thirteen CDGs are closely associated with the sperm motility (SP) signalling pathway with −log p = 19.6 (Fig. 4A). Among them 67 (59%) are CDPKGs (Supplementary Table S9 and S10). Among the CDPKGs involved in SP signalling, a series of protein kinases showed high fold-enrichment (FEG>100), e.g. PRKAG2: 251, PRKCE: 161, PRKCB: 133, PRKCZ: 115, and PRKAR1B: 112, which indicates that protein kinases play crucial roles in the activity of cisplatin, meriting further investigation as a potential targets for cancer treatment.

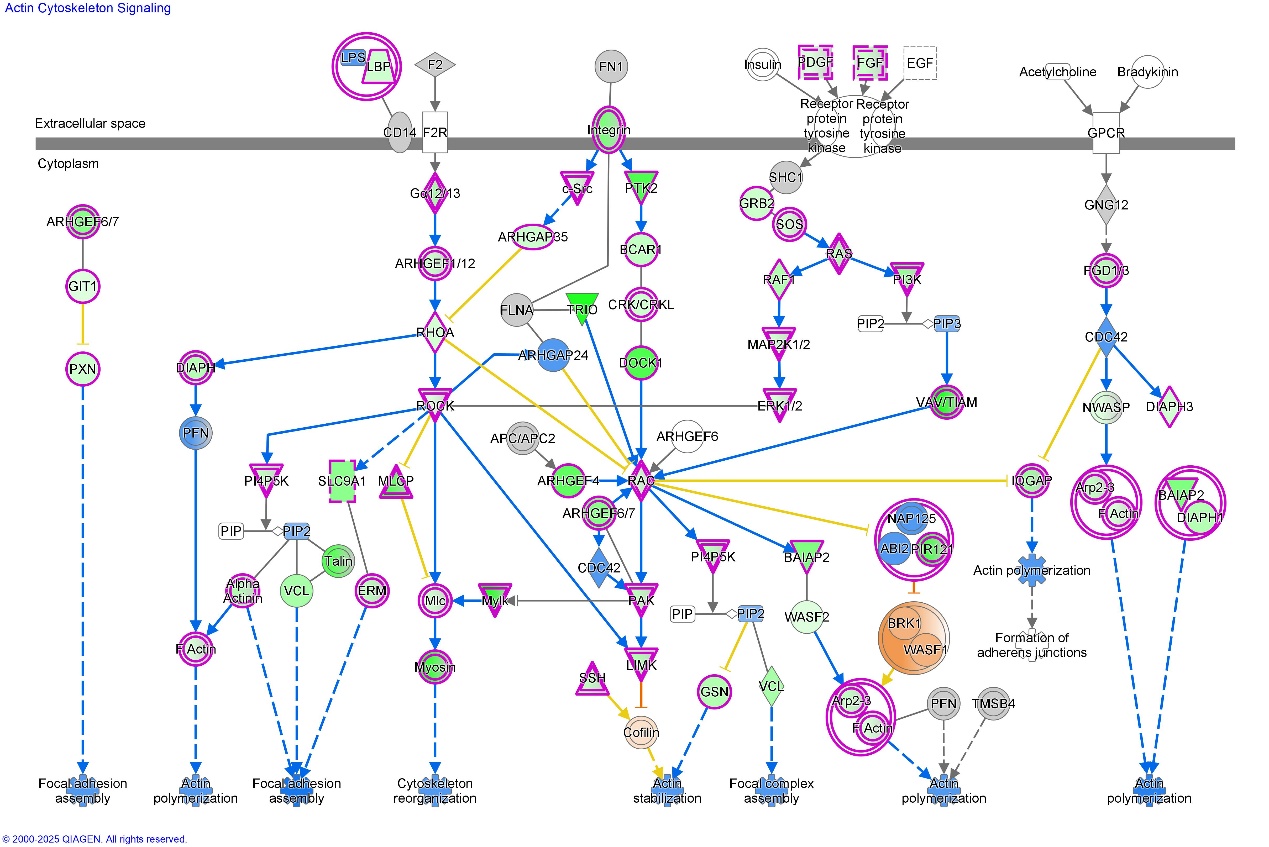

Supplementary Fig. S10. Actin Cytoskeleton Signalling Pathway.

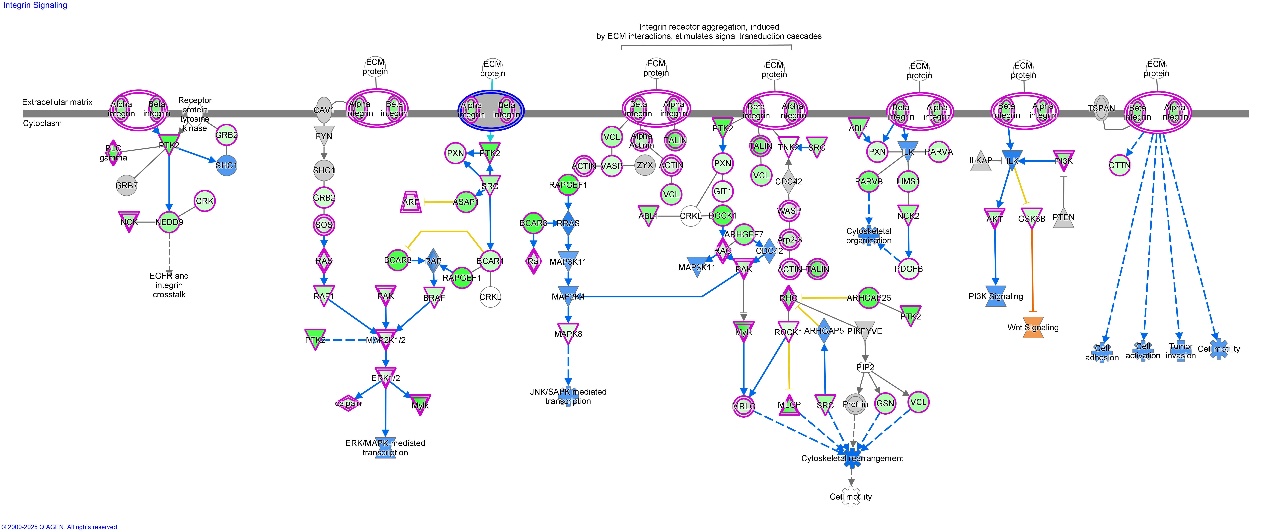

Supplementary Fig. S11. Integrin Signalling Pathway.

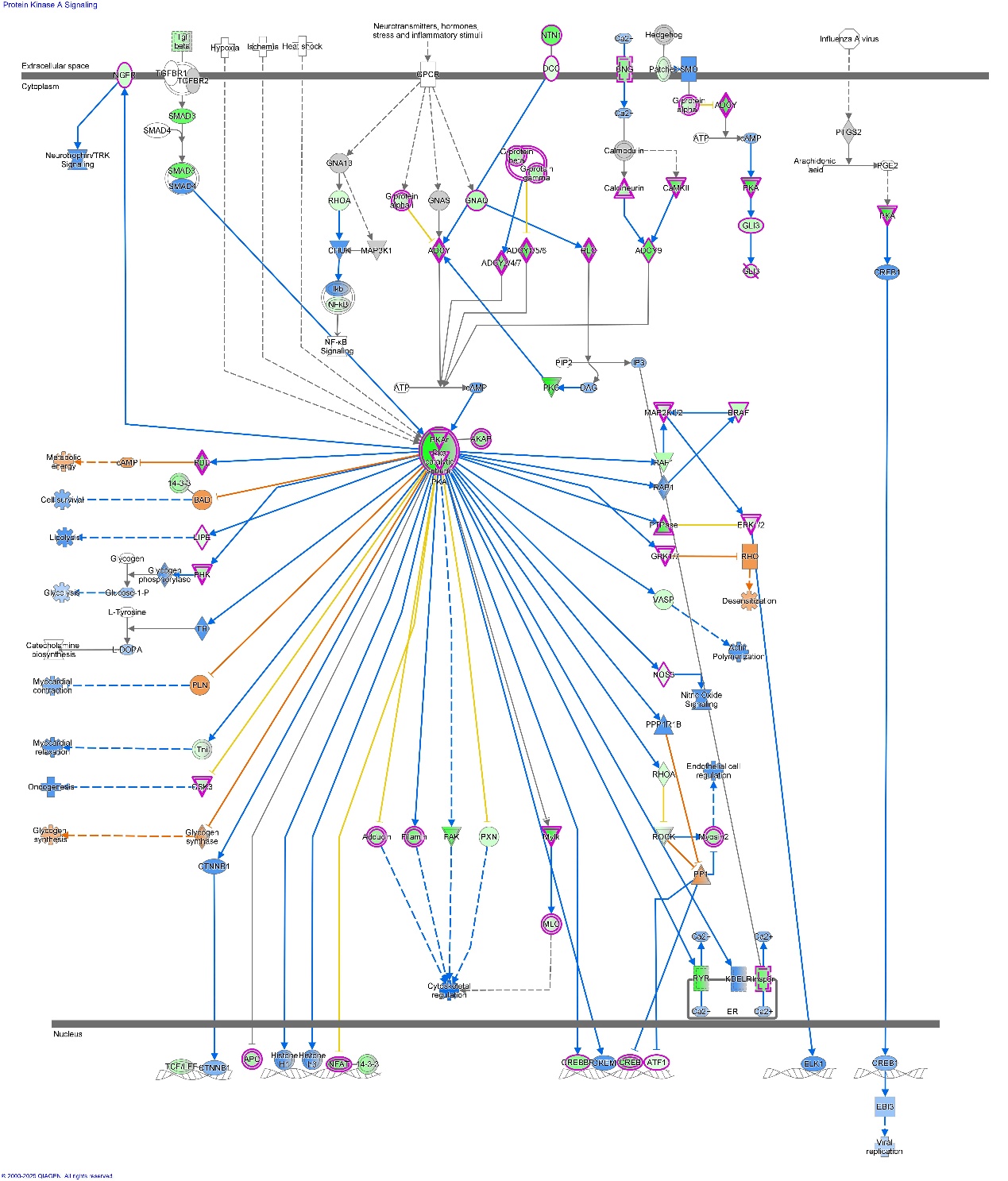

Supplementary Fig. S12. Protein Kinase A Signalling Pathway.

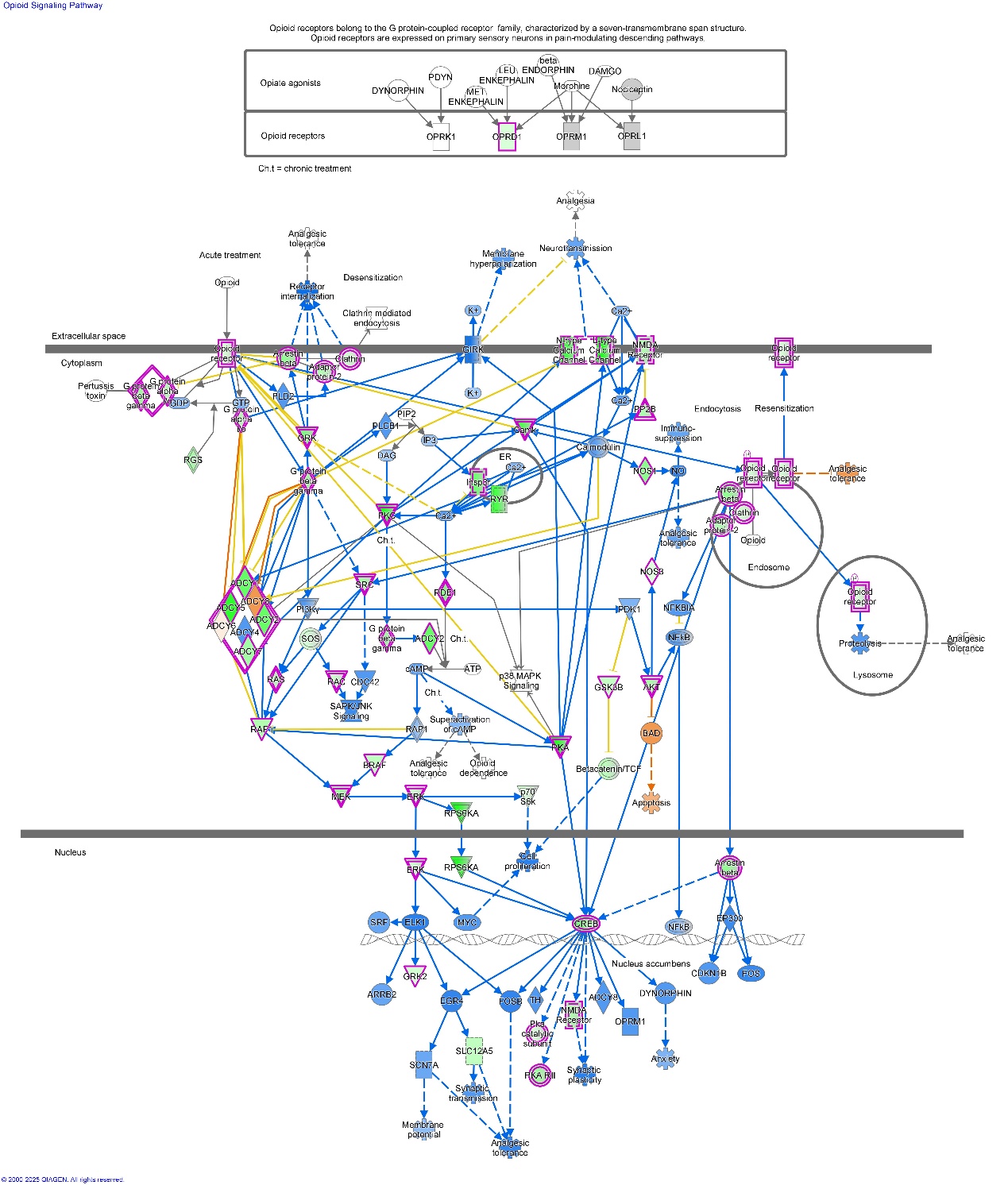

Supplementary Fig. S13. Opioid Signalling Pathway.

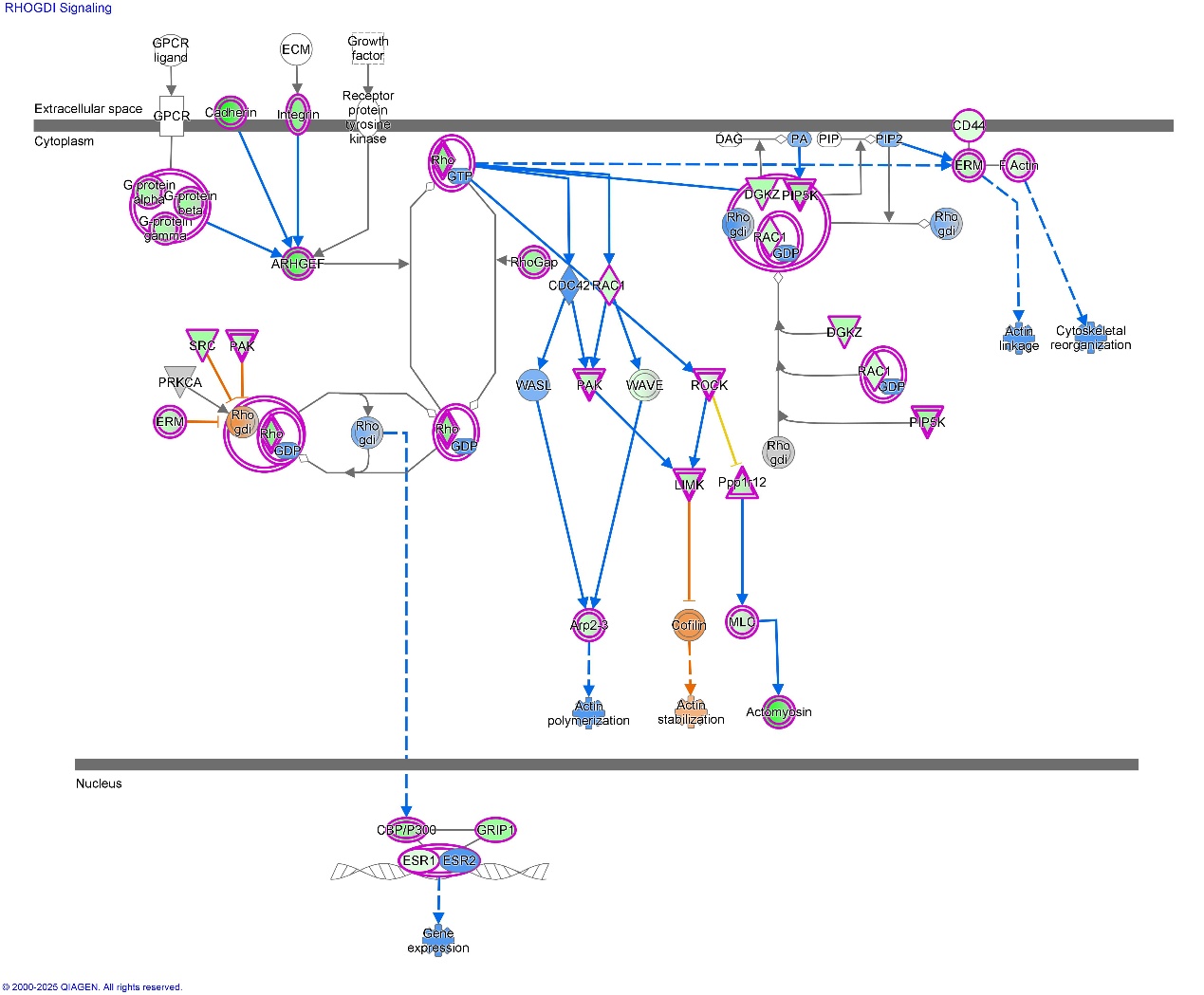

Supplementary Fig. S14. RHOGDI Signalling Pathway.

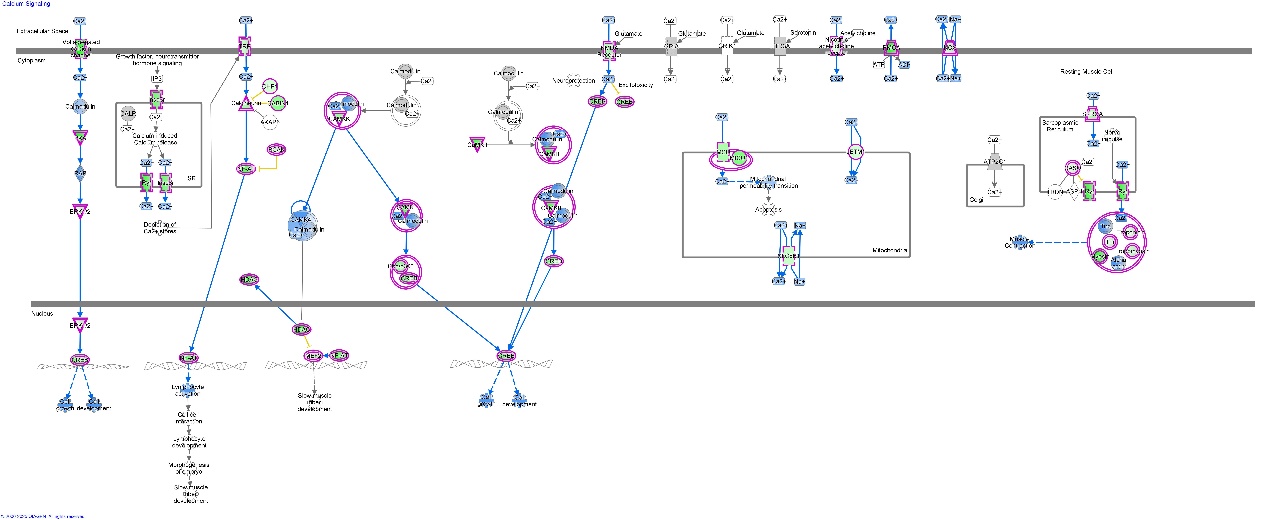

Supplementary Fig. S15. Calcium Signalling Pathway.

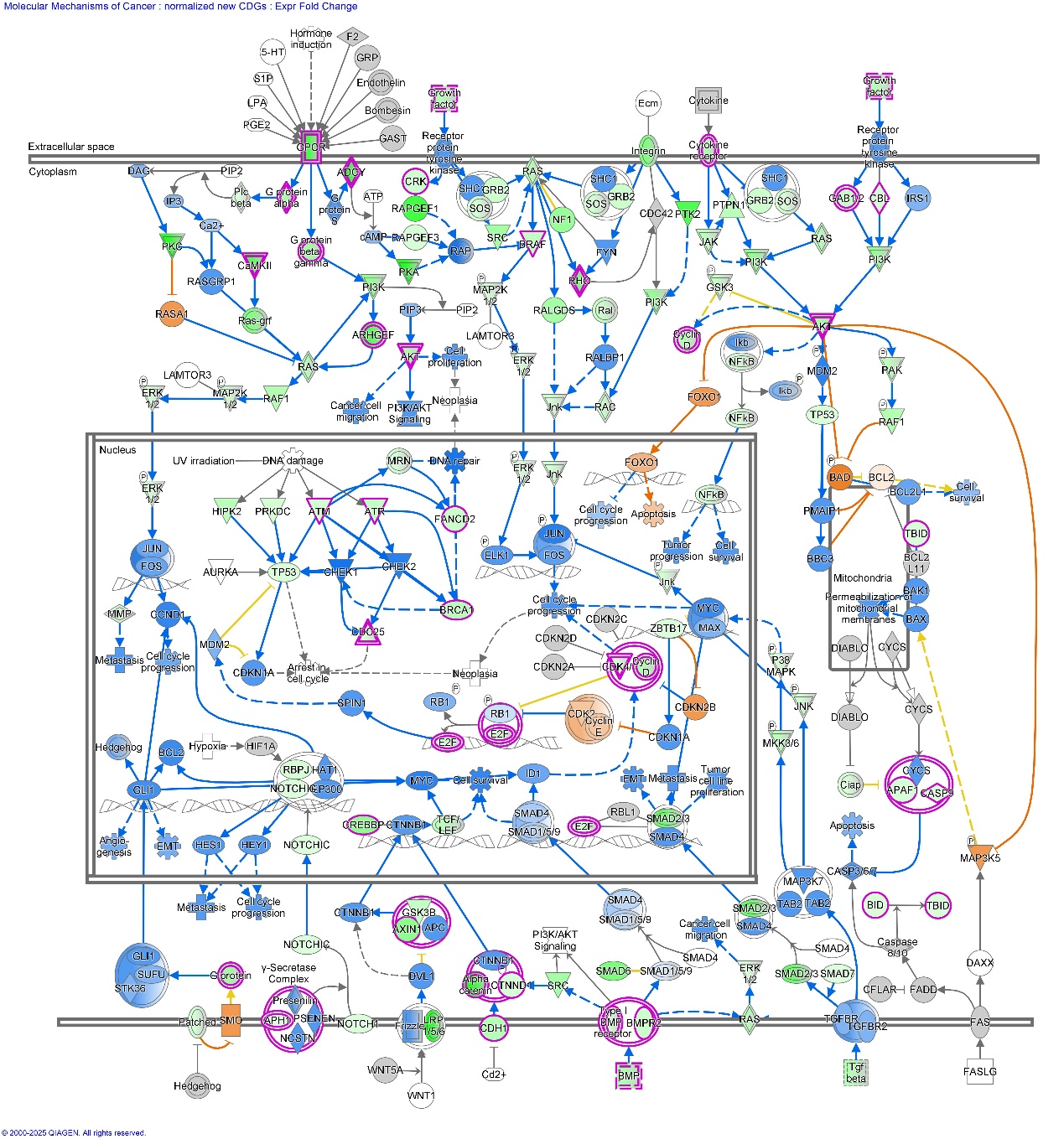

Supplementary Fig. S16. Molecular mechanisms of cancer signalling pathway.

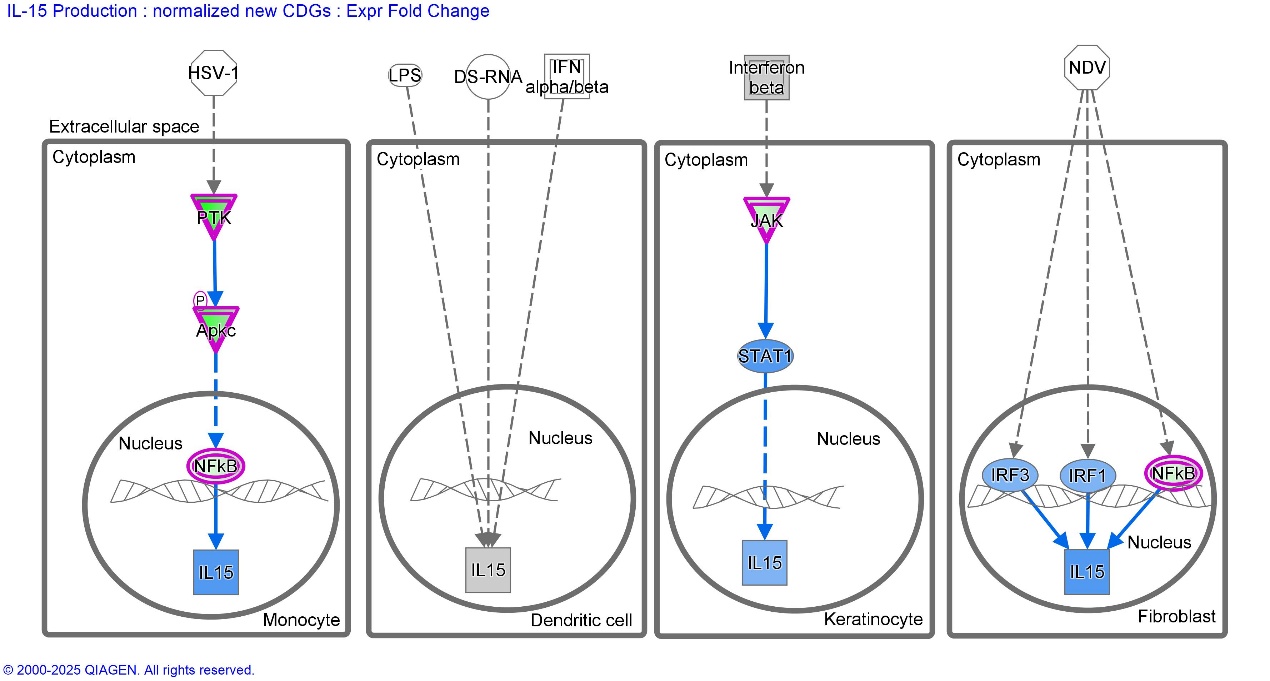

Supplementary Fig. S17. IL-15 production signalling pathway.

Interestingly, bioinformatics analysis showed that 57 CDPKGs cover 95.0% of all CDGs involved in the IL-15 production signalling pathway (−log *p* = 12.7) and are predicted to inhibit IL-15 production with a *z* score of −7.7 (Supplementary Table S9). IL-15 production is activated by PTK-PKC phosphorylation signalling, which induces the activation of JAK kinases, and phosphorylates and activates transcription activators STAT3, STAT5 and STAT6, which may promote the expression of apoptosis inhibitor BCL2L1/BCL-x(L), thus hampering apoptosis[14]. These imply that cisplatin damage to protein kinase genes of the IL-related signalling pathways may play important roles in the activity of cisplatin, e.g., inducing apoptosis by inhibiting the expression of apoptosis inhibitors such as BCL2L1/BCL-x(L).

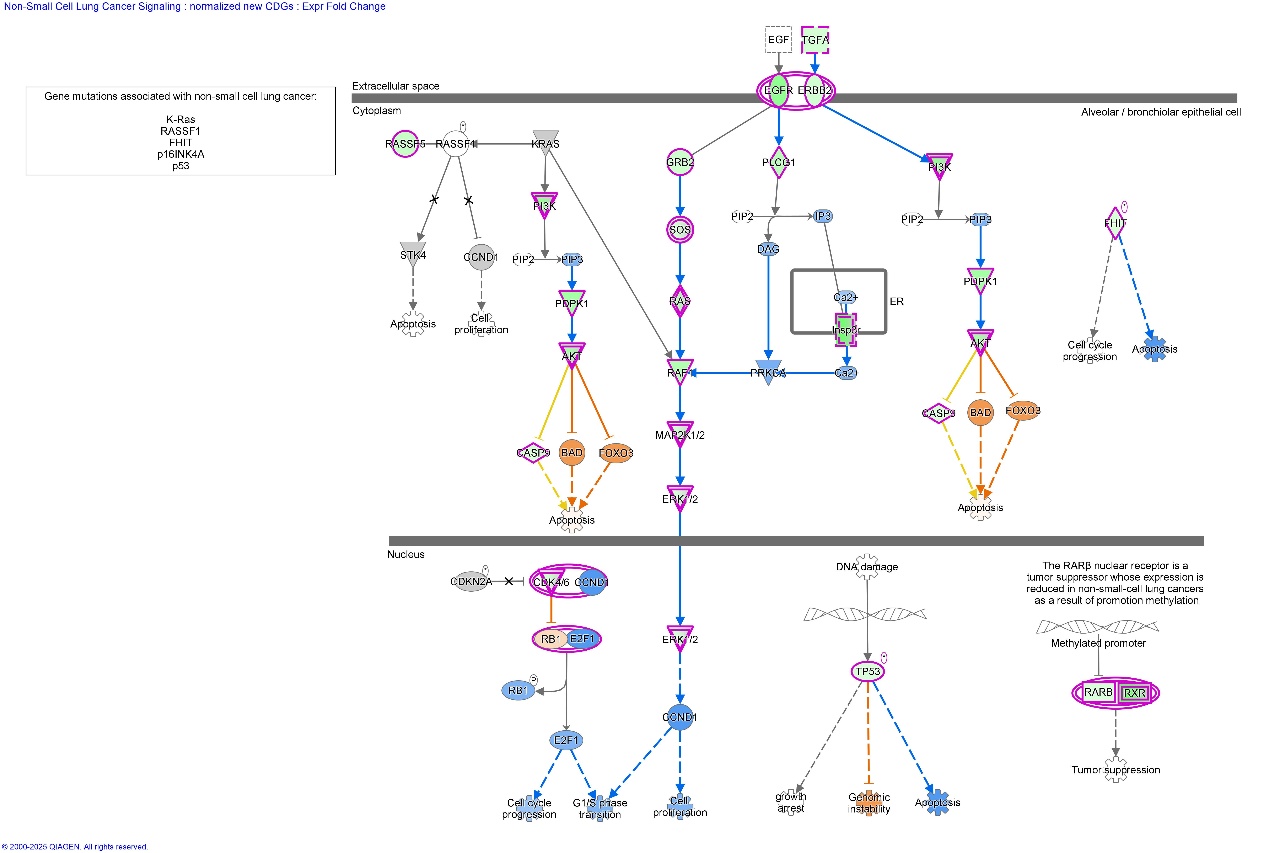

Supplementary Fig. S18. Non-small cell Lung cancer signalling pathway.

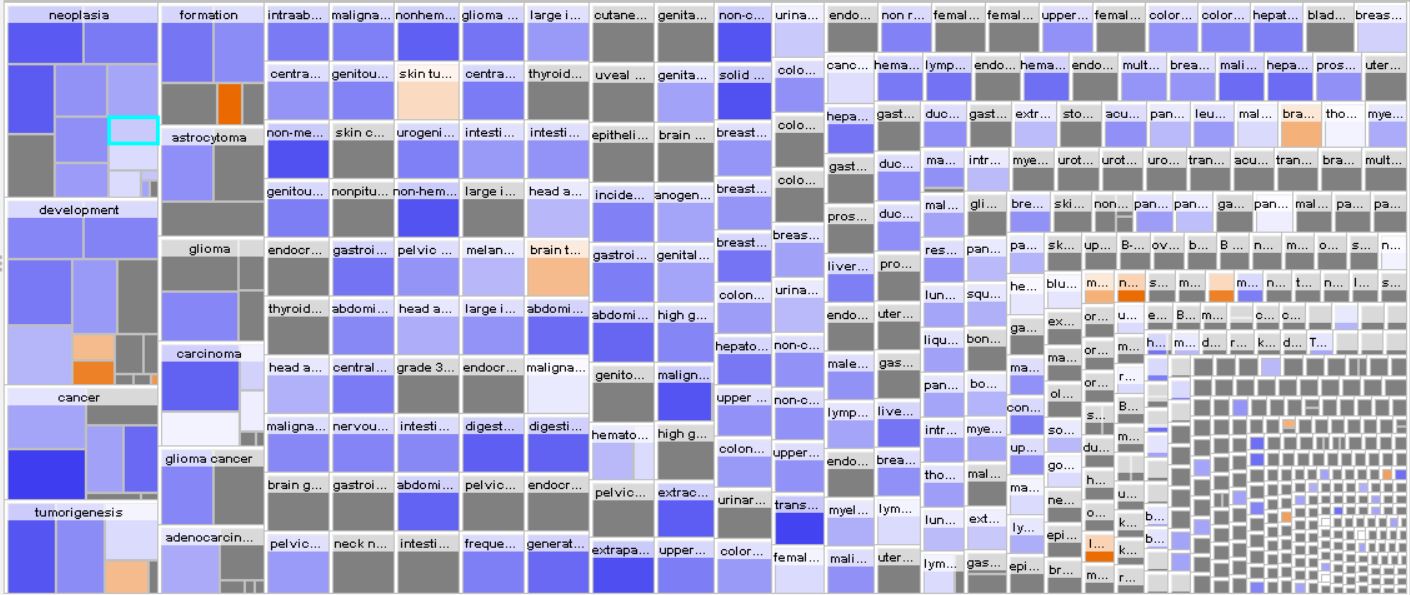

Supplementary Fig. S19. Highly related diseases and bio functions-cancer.

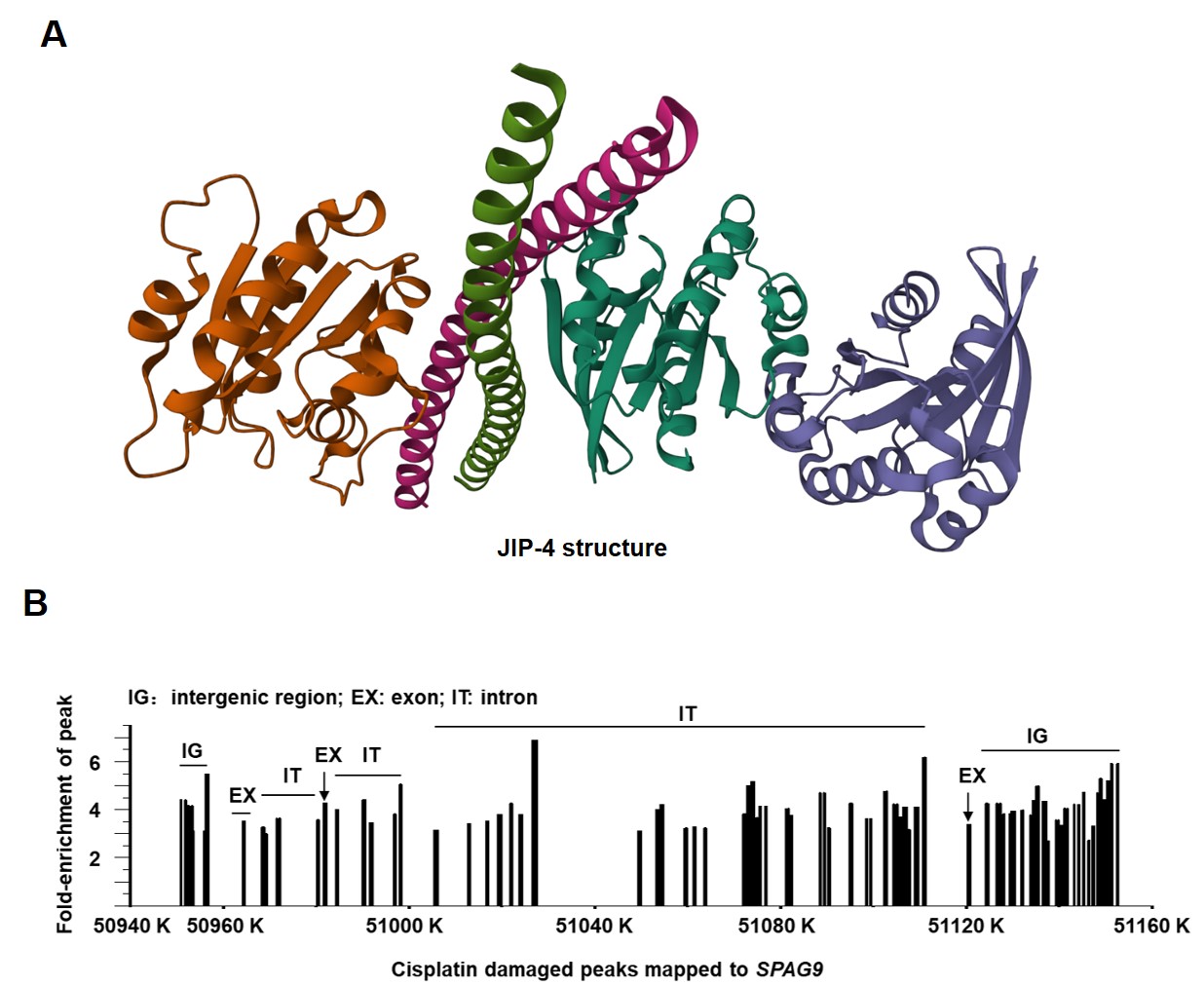

Supplementary Fig. S20. JIP-4 protein and Cisplatin damaged peaks mapped to SPAG9 signalling pathway. (A) Crystal structure of C-jun-amino-terminal kinase-interacting protein 4 (JIP-4) encoded by *SPAG9*, showing four subunits (PDB ID: 2W83). (B) Map of all cisplatin-damaged gene sequences in *SPAG9*, the x-axis numbers show the location of *SPAG9* in chromosome 17. Only cisplatin damage located in the promotor and exon regions will inhibit the expression of protein encoded by the gene.

The expression level of a specific gene is greatly influenced by the location of cisplatin damage in the promoter or exon regions. For example, for *SPAG9* (Supplementary Table S16), no cisplatin lesions were mapped on the promotor sequences, and cisplatin damaged the non-encoding regions (introns) much more than the encoding regions of the gene (exons). This may explain why the full-length JIP-4 decreased in abundance, whereas one or two subunits of JIP-4 increased in abundance (Fig. 5A).

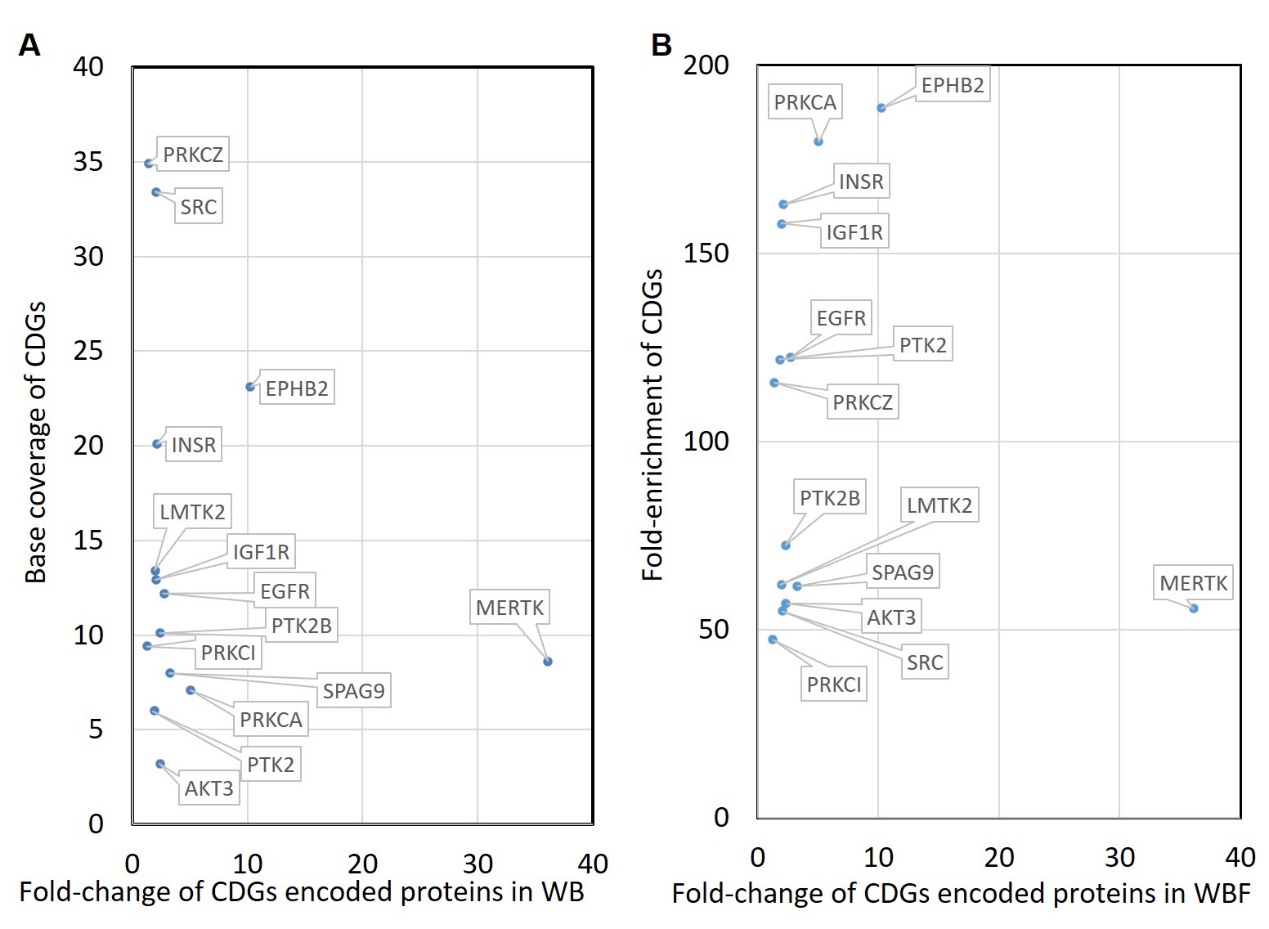

Supplementary Fig. S21. Correlation plots of the fold-change in expression of 14 proteins determined by Western Blot (WB) assays with (A) the fold-enrichment (FE_G_), and (B) the base coverage of the CDGs encoding the 14 proteins.

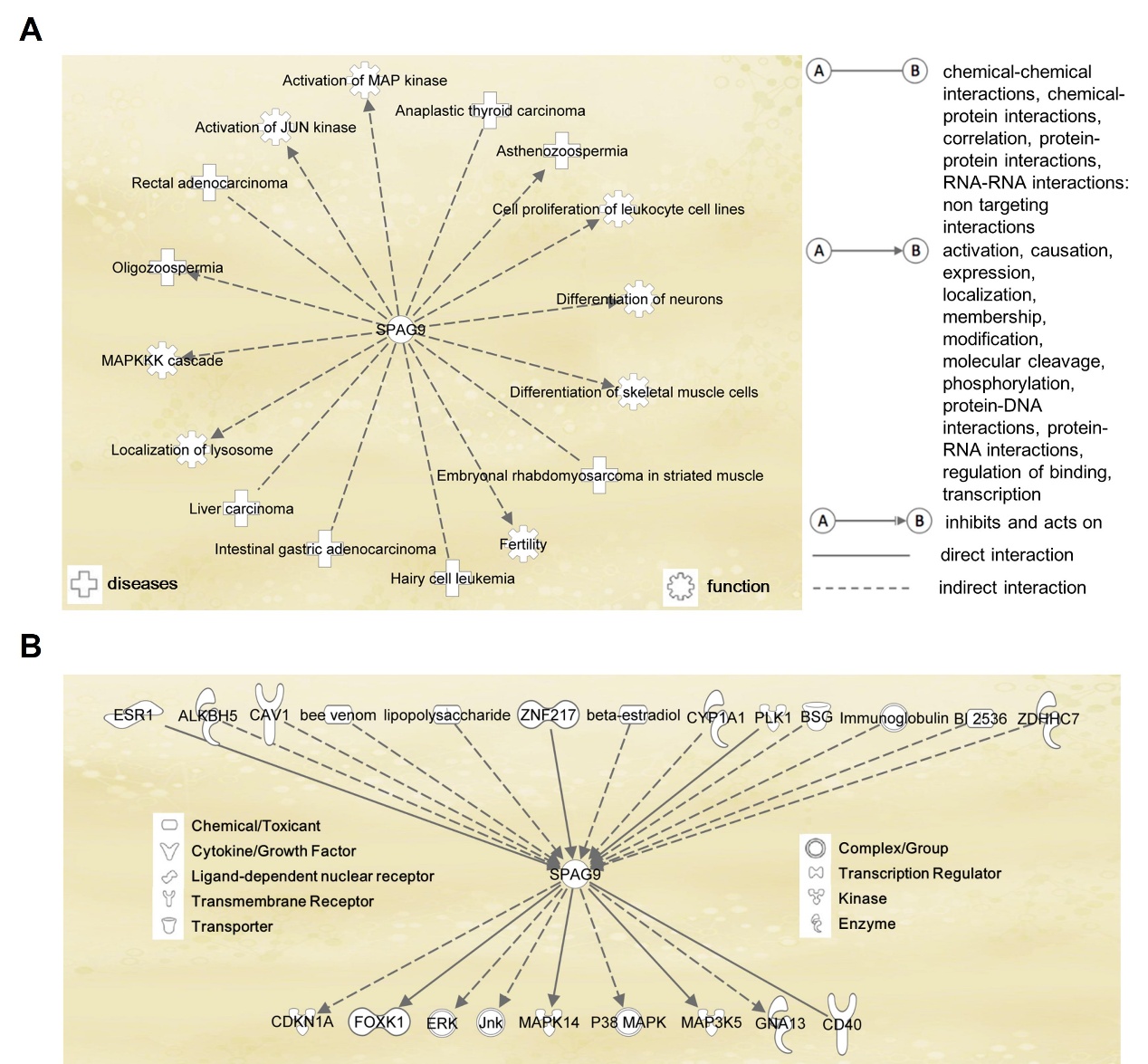

Supplementary Fig. S22. The key roles of SPAG9 in (A) biological processes, and (B) mediating phosphorylation signalling pathways, such as MAPK and PI3K.

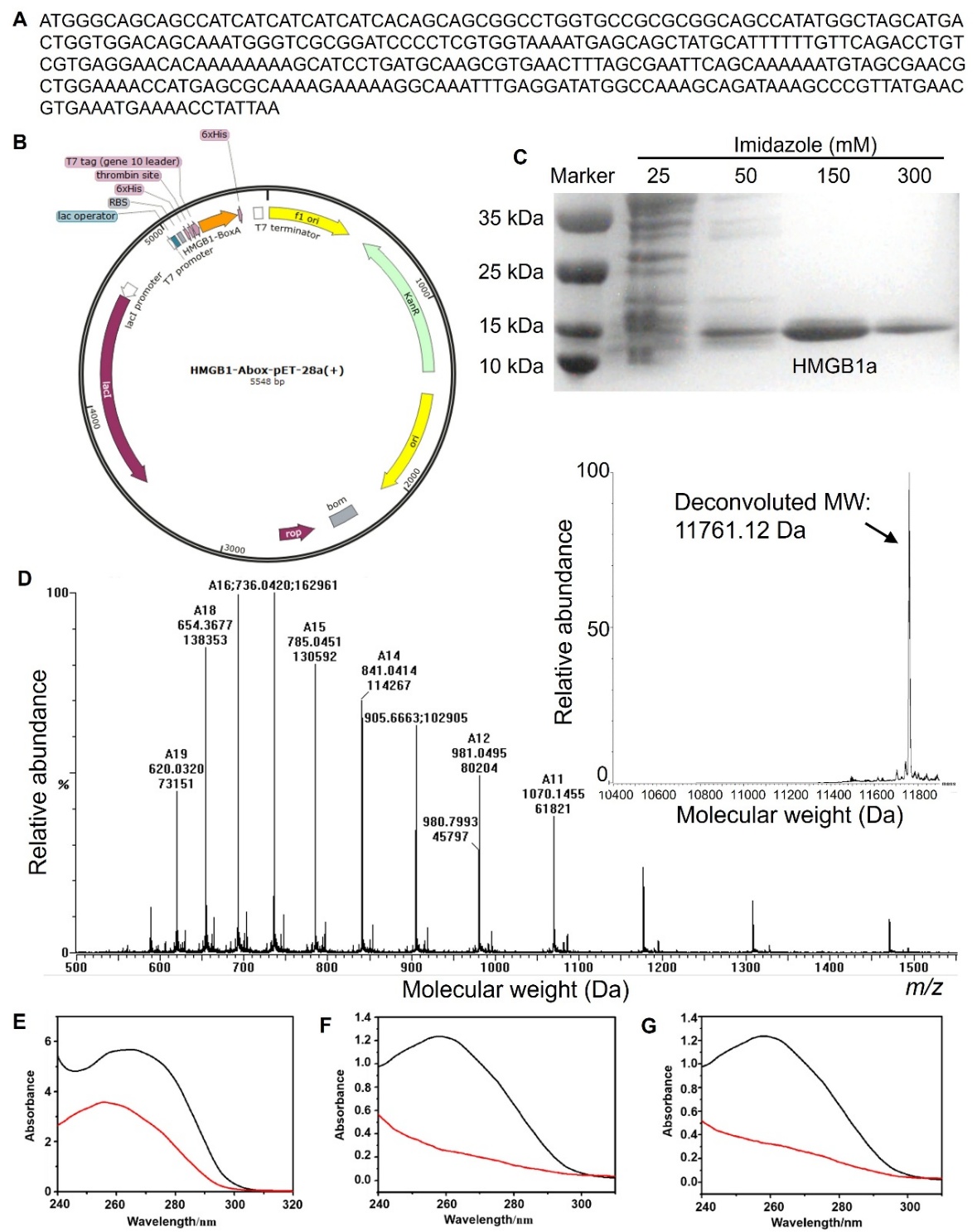

Supplementary Fig. S23. Identification of HMGB1a expressed in *E. coli*. (A) The cDNA sequence of human HMGB1 domain A (HMGB1a) (B) Diagram of vector for HMGB1a expression in *E. coli*. (C) Coomassie-blue-stained gel of HMGB1a expressed in *E. coli.* (D) Mass spectrum of HMGB1a expressed in *E. coli*. (E) UV absorption spectrum of HMGB1a in the supernatant before (black) and after (red) incubation with Ni-modified magnetic beads suspended in PBS. The significant decrease in absorbance over 240 – 300 nm for the supernatant indicates the loading of HMGB1a onto the Ni-modified magnetic beads. (F – G) UV absorption of the DNA fragments from (F) a positive sample, and (G) a control sample, for the supernatant before (black) and after (red) incubation with HMGB1a-functionlized microprobes. The absorbance at 260 nm of the DNA fragments from both positive and control samples significantly decreased after incubation with the affinity microprobe.
